# Humanized nucleosomes reshape replication initiation and rDNA/nucleolar integrity in yeast

**DOI:** 10.1101/2023.05.06.539710

**Authors:** Luciana Lazar-Stefanita, Max A. B. Haase, Jef D. Boeke

## Abstract

Eukaryotic DNA wraps around histone octamers forming nucleosomes, which modulate genome function by defining chromatin environments with distinct accessibility. These well-conserved properties allowed “humanization” of the nucleosome core particle (NCP) in *Saccharomyces cerevisiae* at high fitness costs. Here we studied nucleosome-humanized yeast-genomes to understand how species-specific chromatin affects nuclear organization and function. We found a size increase in human-NCP, linked to shorter free linker DNA, supporting decreased chromatin accessibility. 3-D humanized-genome maps showed increased chromatin compaction and defective centromere clustering, correlated with high chromosomal aneuploidy rate. Site-specific chromatin alterations were associated with lack of initiation of early origins of replication and dysregulation of the ribosomal (rDNA and rRNA) metabolism. This latter led to nucleolar fragmentation and rDNA-array instability, through a non-coding RNA dependent mechanism, leading to its extraordinary, but entirely reversible, intra-chromosomal expansion. Overall, our results reveal species-specific properties of the NCP that define epigenome function across vast evolutionary distances.

**Highlights:** Humanized nucleosomes wrap 10 additional nucleotides, shortening free linker length

Histone-humanized nucleosomes have increased occupancy for DNA

Humanized nucleosomes potentially decrease chromatin accessibility by blocking-out free linker DNA

Nucleosome humanization impedes DNA replication by affecting chromatin structure at origins

Humanized nucleosomes reversibly destabilize the ribosomal DNA array and leads to massive intrachromosomal rDNA locus expansion

Histone humanization disrupts rDNA silencing and leads to nucleolar fragmentation

## Introduction

In eukaryotes DNA molecules are packed in the nucleus in a hierarchical folding structure. The first level of organization consists of ∼1.7 superhelical turns of 147 bp DNA around an octamer of two copies each of the four histone proteins: H2A, H2B, H3 and H4 (reviewed in McGhee and Felsenfeld^1, 2^). Altogether they form the nucleosome core particle (NCP) that constitutes the basic structural unit of chromatin^2, 3^, conserved throughout eukaryotes (reviewed in Kornberg and Lorch^4^, Malik and Henikoff^5^). Aside from the canonical core histones, the sequence-specified centromere of *Saccharomyces cerevisiae* (∼120 bp AT-rich region) is organized into a specialized nucleosome containing Cse4, a centromere-specific variant of histone H3^6–8^. A single Cse4 nucleosome is thought to form the minimal unit of the point centromeric chromatin^7, 9^, required for the recruitment of the kinetochore complex and proper chromosome segregation (reviewed in Cleveland et al.^10^). Genome-wide footprinting showed that the centromeric nucleosome contains a micrococcal nuclease-resistant unit of ∼123–135 bp, significantly shorter than the canonical nucleosome^11^.

Nearly 80% of the yeast DNA is incorporated into stable nucleosomes^12^. The “chromatinization” process takes place primarily during S-phase and is coupled to the passage of the DNA replication fork^13^, when parental and de novo synthesized histones are deposited onto the two nascent DNA molecules^14–17^. During this process, histone chaperones and nucleosome remodelers closely interact with components of the replication machinery to deposit new and old histone octamers onto the newly replicated duplexes^18^ (reviewed in Budhavarapu et al.^19^, Sauer et al.^20^). The DNA sequence interconnecting consecutive NCPs to form higher-order structures is called the linker^4^. Its length varies among cell types and organisms (e.g., ∼20 bp long in *S. cerevisiae*^21^), and is thought to be inversely correlated with gene activity^22–25^. In addition, correctly stabilizing nucleosome positions on the DNA polymer - relative to *cis* regulatory elements (e.g., replication origins and transcription start sites) - is a critical component of genome function and regulation (reviewed in Rando and Chang^26^, Lai and Pugh^27^). Regions of the genome that are devoid of nucleosomes, referred to as nucleosome-free regions (NFRs), represent accessible parts of the chromatin where multiprotein complexes can assemble and regulate/perform key DNA-templated processes e.g., replication and transcription^27^. As these processes occur in the context of the surrounding chromatin environment, nucleosome occupancy and positioning can restrict access to particular DNA sequences and influence genome-wide initiation/firing of origins of replication^28, 29^ and transcription regulation (reviewed in Bai and Morozov^30^). Ergo post-translational modifications of nucleosomes, defining distinct local chromatin environments, also affect these processes (reviewed in Bowman and Poirier^31, 32^). Furthermore, it has been proposed that the epigenetic information is maintained during transcription and replication^33^, and can even be transmitted through generations (reviewed in Kaufman and Rando^34^, Campos et al.^35^). On the other hand, yeast nucleosomes that lack certain post-translational modifications (e.g., acetylation) are associated with silenced chromatin. The hypoacetylated nucleosomes promote a compact chromatin structure (or heterochromatin) and makes DNA inaccessible to processes such as transcription and replication initiation (reviewed in Gartenberg and Smith^36^). The Sir2 (Silent information regulator 2) protein is a conserved NAD^+^-dependent deacetylase that removes key acetyl groups from histone H3 (H3K9 and H3K14) and H4 (H4K16)^37, 38^. This type of epigenetic silencing occurs at diverse genomic sites including the silent *HM*-mating type loci, telomeres and the ribosomal DNA (rDNA) tandem array. Hypoacetylated H3 and H4 histones^39^ have been linked with heterochromatin formation by establishing direct interactions with components of the silencing complex (Sir3/4) and nucleosomes^40–42^. These interactions not only participate in telomere and *HM* loci silencing, but also in their nuclear envelope positioning^43^. Finally, only Sir2 deacetylase activity is required for rDNA silencing whereas Sir3/4 are dispensable^44, 45^.

Despite the vast evolutionary distance between yeast and human, large-scale systematic studies have found that several hundred yeast genes can be individually replaced with their human orthologs and sustain yeast growth (3%)^46, 47^ (reviewed in Dunham and Flower^48^). Multigene interspecies swaps have also been reported^49^, exemplified by the humanization of the entire NCP (H2A, H2B, H3 and H4) of *Saccharomyces cerevisiae* (Truong and Boeke^50^). Although yeast and human histones are highly conserved (68% - 92% identity), isolation of NCP-humanized yeasts required acquiring genetic mutations to survive and they display dramatic phenotypic defects associated with a global RNA reduction. These findings presented the opportunity to examine the effects of human histones in yeast to provide valuable insights into the mechanisms that govern chromatin-related processes in distantly related organisms. Here we used yeast strains that rely on human histones for packaging their DNA molecules, to address how these species-specific units of chromatin structure alter yeast genome organization - from nucleosome fibers up to the 3D structure of chromosomes - and how these structural changes reflect upon biological processes (e.g., DNA replication, gene silencing and genome stability).

## Results

To ascribe our findings to the type of histone used for DNA packaging (yeast vs. human), we compared results obtained from humanized strains carrying distinct “humanization-suppressing” mutations. This terminology was coined to indicate a specific subset of genetic mutations required for histone-humanized yeasts to survive and propagate^50^. Notably, some of these mutations were found to have distinct effects on genome stability, leading to isolation of two classes of humanized strains with distinct levels of aneuploidy. Here we focused on two such humanized yeast strains carrying each a single point mutation, in either *DAD1* (strain: yDT180, *dad1*-E50D) or *SCC4* (strain: yDT92, *scc4*-D65Y) genes, which display either normal or abnormal ploidy, respectively. These were isolated by Truong in 2017, and the mechanism of ploidy stabilization was addressed by Haase et al.^51^ who documented that the mutation in *dad1*- E50D (component of the outer kinetochore DASH/Dam1 complex) stabilized ploidy of the histone-humanized yeasts by weakening the interaction between the outer kinetochore and the microtubules.

### Visualization of histone-humanized chromatin fibers in yeast

Previously reported nucleosome occupancy maps have shown a high degree of structural conservation of histone-humanized chromatin fibers in yeast^50, 51^, with notable exceptions described later in this manuscript. Even if nucleosome positioning appears to be well conserved overall, the details of the wrapping of yeast DNA on the human NCP remain unknown. To address this question, we used transmission electron microscopy (TEM) to directly image chromatin fibers extracted from yeast cells harboring either human or native yeast histones. Representative images in Figure 1A show the expected “beads-on-a-string” array arrangement of the nucleosomes in the wild-type*, Sc* (*Saccharomyces cerevisiae*), and in the histone-humanized, *Hs* (*Homo sapiens*), cells. The schematic in Figure 1B shows an example of an NCP used as a benchmark to calculate mononucleosome surface area. The latter was measured using chromatin images acquired at various resolutions (representative images are shown in Figure S1 and S2) accounting for approximately 500 nucleosomes for each *Sc* and *Hs* strain (Table S1). Bee swarm plots in Figure 1C point to a small but significant increase in the surface of the mononucleosome in both histone-humanized strains (*Hs*: yDT92 and yDT180), relative to the WT (*Sc*: BY4742) yeast strain. This change corresponds to a ∼5% increase in the circumference of the NCP (∼1.8 nm) and suggests that more DNA in the nucleosome repeat length is wrapped/protected by the histone-humanized octamer compared to the native yeast one. Moreover, this result is quantitatively supported by an orthogonal method that directly measured DNA fragment size (using capillary electrophoresis) from MNase digested chromatin and showed that the *Hs-*NCP protects ∼10 bp more DNA than the *Sc-*NCP (Figure 1D data from Haase et al., co-submission). Importantly, this result excludes the possibility of imaging and analyzing semi-complete NCPs from yeast due to the lower intrinsic stability of the histone octamer in yeast compared to other metazoans^52, 53^ (reviewed in McGinty and Tan^54^). Since we did not detect any change in nucleosome positioning nor in the repeat length itself, it suggests that the extra 10 bp of protected sequence corresponds to a reduction in free linker DNA length in yeast. This finding provides in vivo reinforcement to the idea that human histone octamers associate more stably with DNA than the yeast ones and nucleosome packaging in higher-order chromatin structures is fundamentally different between these two species^52, 55^. Finally, these results provide a structural underpinning for the general decrease in chromatin accessibility in the humanized yeast and provides a plausible explanation for their global downregulation of RNA (previously reported in Truong and Boeke^50^, Haase et al.^51^), presumably reflecting reduced access of RNA polymerases and/or transcription factors to DNA.

**Figure 1.**
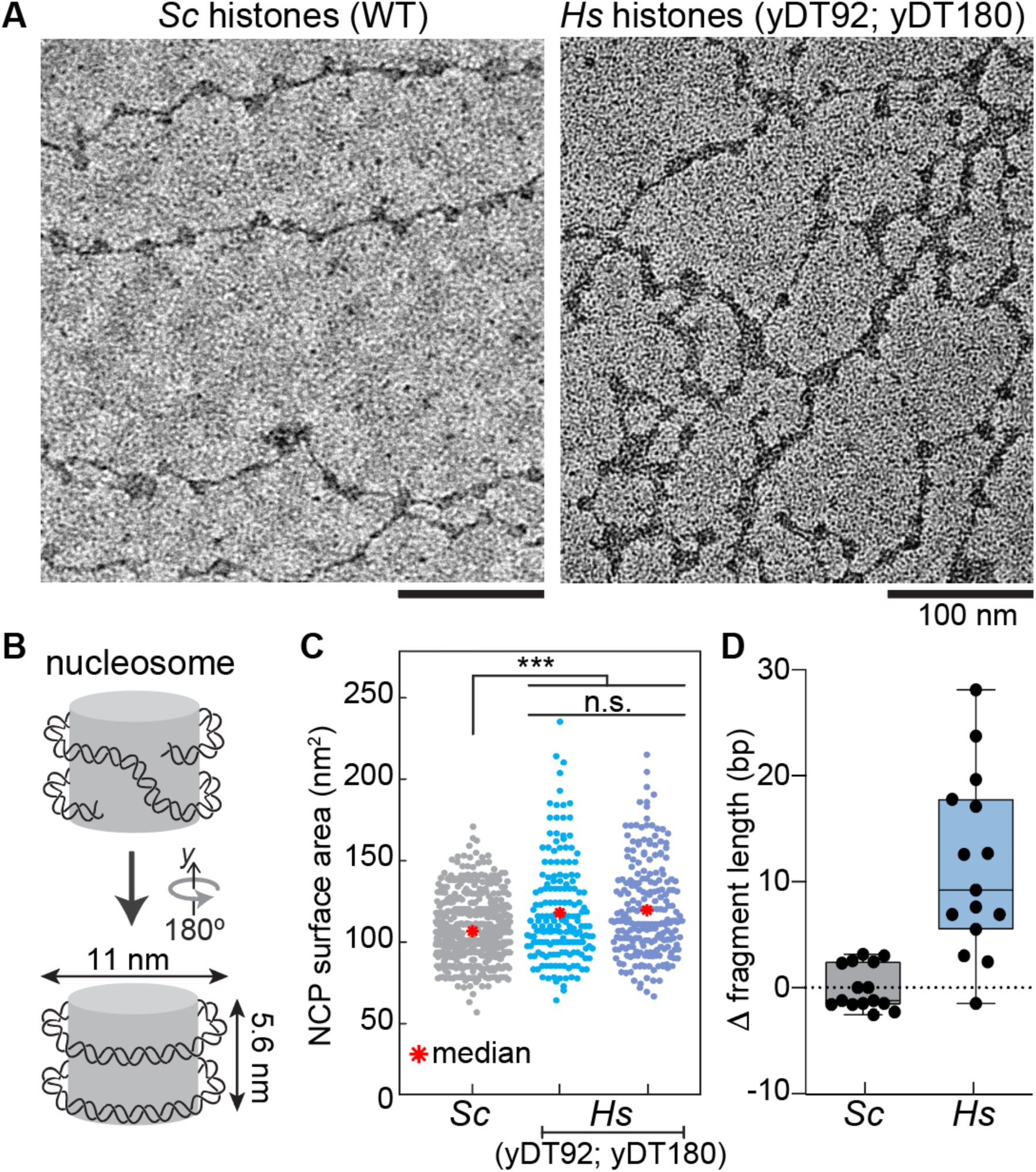
Visualizing histone-humanized nucleosome fibers in yeast. (**A**) Representative electron microscopy images showing 10 nm nucleosome fibers. Left, wild-type yeast with native histones (*Saccharomyces cerevisiae, Sc,* strain: BY4742; see also Figure S1). Right, histone-humanized (*Homo sapiens, Hs*, strains: yDT92, yDT180 fibers; see also Figure S2). (**B**) Schematic representation of the nucleosome core particle (NCP) with dimensions in nm. (**C**) Bee swarm plots showing the average estimated NCP surface area (nm^2^) in the wild-type (*Sc*) and histone-humanized strains (*Hs*). Median, S.D. and *P* values (*** *P* <0.0015; n.s. *P* > 0.05) were calculated using a two tailed t-test function (Table S1). (**D**) Boxplots quantifying the difference of the nucleosome fragment length in *Hs* relative to *Sc* (DNA fragment length analysis of MNase digested chromatin; data from 3 biological replicates: comparisons of lengths from monoup to penta-nucleosome fragments are shown by each dot Haase et al., co-submission).

### 3-Dimensional organization of the histone-humanized yeast genome

Next, we investigated if the nanoscale effects of histone humanization are echoed at longer genomic distances, affecting the overall spatial organization of chromosomes. Genome-wide proximity maps of the *Hs* and *Sc* yeast strains were generated using the chromatin conformation capture approach, Hi-C^56^. At first glance, the 2D (2-Dimensional) interaction/contact frequency maps of two representative chromosomes (chr *IV* and chr *V*) in Figure 2A show that the typical organization of *S. cerevisiae*’s genome is preserved overall in both histone-humanized suppressor mutants (*Hs dad1*-E50D and *Hs scc4*-D65Y). The so-called Rabl-like organization^57^ of yeast chromosomes is characterized by the spatial clustering of all peri-centromeric regions (indicated with black arrowheads in Figure 2A) and their relative insulation from the chromosomal arm sequences (Figure 3C, upper schematic)^58^. Further analysis of intra-chromosomal contacts, that computes the decay in contact probability (*p*) as a function of the genomic distance (*s*), showed a small but reproducible increase of contacts at mid-range (∼20-50 kb) distances in the *Hs* chromosomes relative to *Sc* (Figure 2B). In addition, the contact variation maps in Figure 2C not only confirmed an increase in mid-range intrachromosomal contacts in the *Hs* strains, as shown by the red signal running parallel to the proximal diagonal, but they also revealed local contact variations surrounding the pericentromeres. Here, the black arrowheads point to peri-centromeric positions which appear to favor interactions with the distal chromosomal arms, both in *cis* (within the same chromosome) and in *trans* (on different chromosomes), in the *Hs* relative to *Sc* maps. These stand out as red contact stripes on the comparison maps and support the hypothesis that the centromeres are de-clustered in the *Hs* strains relative to the *Sc* strain. To test this hypothesis, we used the normalized *Hs* and *Sc* contact maps (expanded versions of the insets shown in Figure 2A) to quantify the frequency of contacts that each peri-centromeric region (50 kb sequence centered on a given centromere) makes with the remaining 15 peri-centromeres, where higher contact values correspond to robust centromere clustering. The left plot in Figure 3A reveals a significant reduction to the inter-centromere contacts in *Hs* compared to *Sc* of approximately 30%. Notably this result was reproduced in both *Hs dad1*-E50D and *Hs scc4*-D65Y yeasts, suggesting that centromere de-clustering occurs regardless of the humanization-suppressor mutations in the histone-humanized strains (Figure 3B, average 3-Dimensional representations of the Hi-C maps with a viewpoint on the centromeres in yellow). Based on DNA content analysis and chromosome coverage plots (Figure S3A-B), we confirmed that a specific subset of chromosomes tended to be aneuploidy in *Hs scc4*-D65Y, whereas the genome of *Hs dad1*-E50D maintains normal ploidy (as previously reported in Truong and Boeke^50^ and Haase et al.^51^). Given that centromeres are the key elements responsible for chromosome stability during cell division, we reasoned that peri-centromeres of aneuploid chromosomes may fail to achieve this function due to a further aggravated defect in their clustering. To explore this hypothesis, we first computed the inter-chromosome contact variations for both *Hs* strains with normal and aneuploid chromosomes (ratio of the normalized Hi-C maps: humanized vs. WT yeast, Figure S3C), which we used as a ploidy-correction to the inter-centromere contact variation between humanized and WT strains. The plot on the right in Figure 3A shows a further reduction (up to ∼45%) in centromere clustering exclusive to the aneuploid (amber) chromosomes in *Hs scc4*-D65Y relative to the euploid (gray) chromosomes. Overall, our results show that humanization of the canonical histones in yeast destabilizes the structure of the peri-centromeric chromatin and leads to centromere de-clustering (Figure 3C, lower schematic). This provides structural support explaining the frequent chromosomal aneuploidies observed post-humanization.

**Figure 2.**
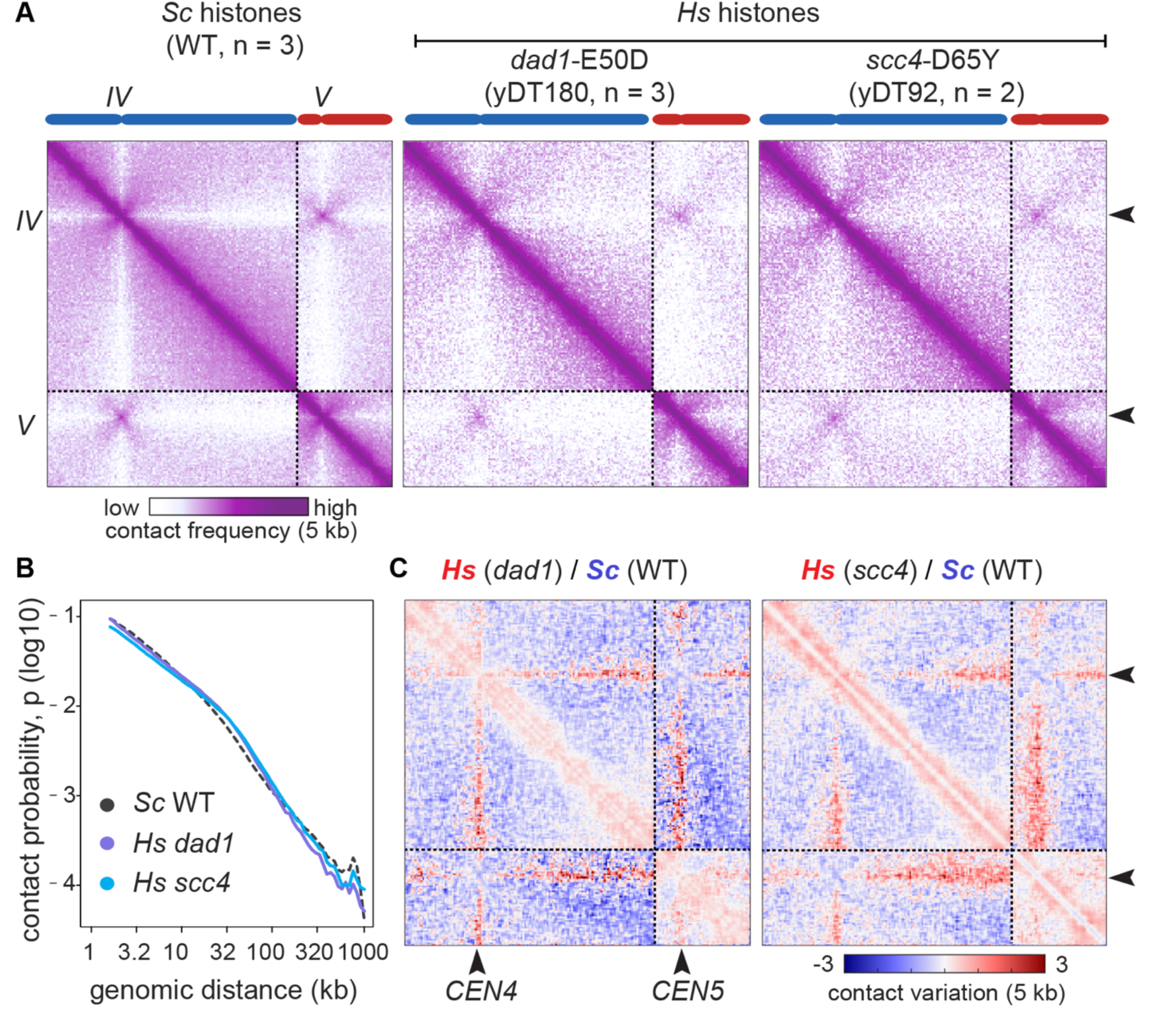
3D genome organization of histone-humanized chromatin. (**A**) Insets of Hi-C contact frequency maps showing chromosome *IV* and *V* underlined by dotted lines in yeast strains with *Sc* histones vs. *Hs* histones carrying distinct humanization-suppressor mutations (yDT180 w. *dad1*-E50D and yDT92 w. *scc4*-D65Y). Blue (*IV*) and red (*V*) chromosomes are plotted on the *x* and *y* axis of the maps binned at 5 kb size resolution. Black arrowheads point at centromere positions, i.e., *CEN4* and *CEN5*. Purple to white color scale indicates increase in contact frequency (log10). (**B**) Contact probability (p) in function of the genomic distance (kb) represents the average decay of the intra-chromosomal contact frequency between loci with the increment in their genomic distances. Replicates of the strains in A were plotted together. (**C**) Comparisons of contact maps in panel A. Log2-ratio maps of each of the *Hs* strains vs. the *Sc* strain. Color bar indicates contact variation between samples (log2 ratio 5 kb-binned).

**Figure 3.**
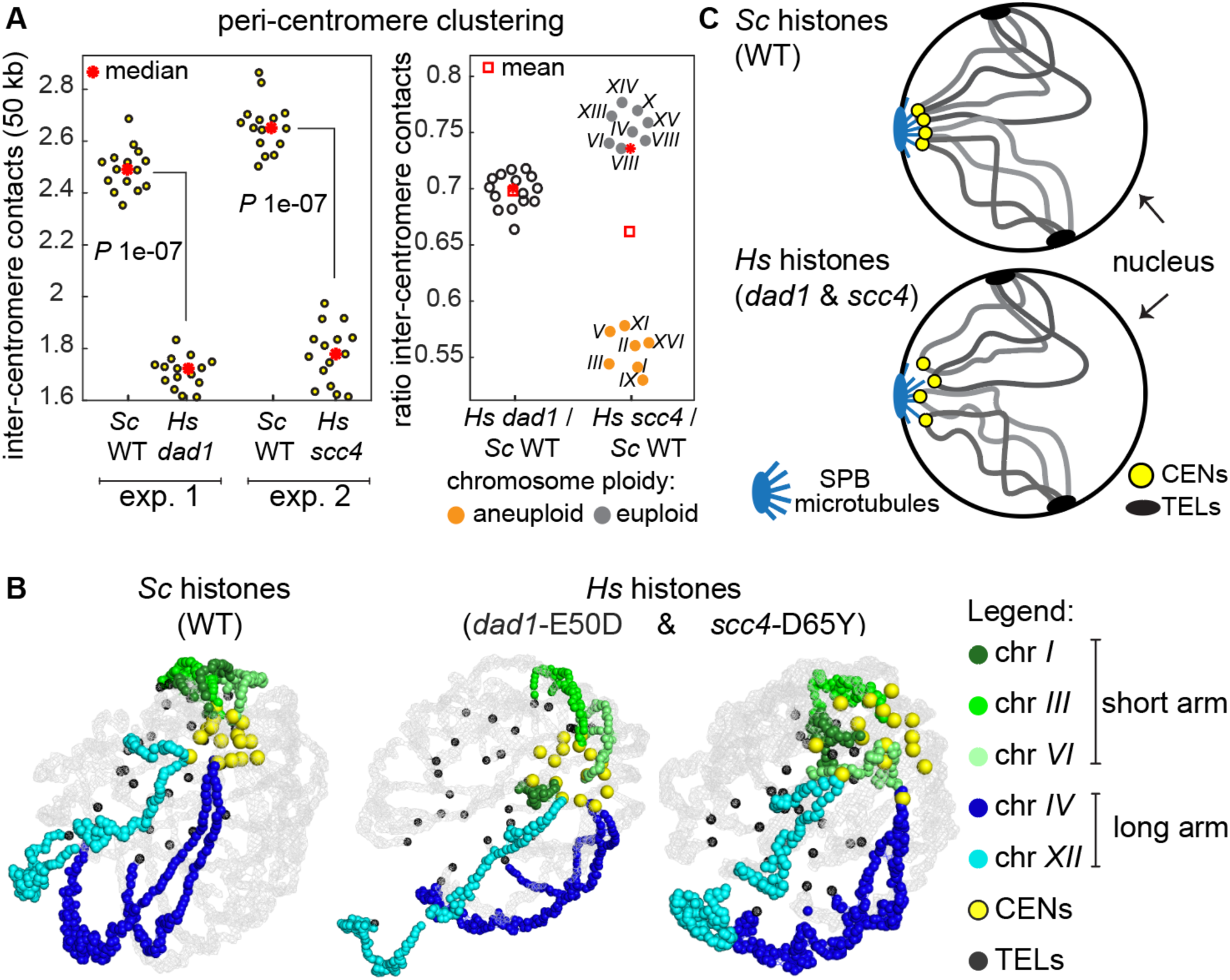
Histone humanization leads to de-clustering of yeast centromeres. (**A**) Centromere clustering in histone-humanized vs. wild-type yeast using normalized Hi-C genome maps. Left plots: quantifications of all inter-centromere contacts, plotted in 50 kb-windows centered on a given centromere (each dot represents the sum of all *trans* contacts a peri-centromeric region makes with the other 15 peri-centromeres) in the *Hs* (yDT180 *dad1*-E50D and yDT92 *scc4*-D65Y) strains relative to the corresponding *Sc* from the same experiment (indicated as exp. 1 and 2). Right plot: variations of inter-centromere contacts in *Hs* vs. *Sc* plotted according to level of chromosome ploidy (aneuploid vs. euploid shown in Figure S3B). (**B**) 3D average representations of the *Sc* and *Hs* corresponding to complete chromosome-contact maps from Figure 2A. Color code highlight a few chromosomes with either short or long arms, as well as centromeres (CENs) and telomeres (TELs). (**C**) Schematic model of Rabl-like organizations of wild-type yeast chromosomes (*Sc* top panel) compared to the histone-humanized (*Hs* bottom panel) one, showing de-clustering of centromeres. Examples of chromosome arms (gray lines) anchored at the nuclear membrane through CENs and TELs.

### Histone humanization delays activation timing of DNA replication origins

Given the inseparable relationship between genome structure and function, we then asked whether structural changes introduced by humanized nucleosomes would affect specific biological processes. Our previous work has shown that the humanized yeasts have low fitness^50^, suffering from a prolonged cell cycle (∼3-fold longer). One potential explanation for the cell cycle delay might be a defect in DNA replication initiation of histone-humanized chromosomes, since recognition of replication origins might be blocked by the increased nucleosome stability/binding.

In *S. cerevisiae*, DNA replication start sites or origins (named Autonomously Replicating Sequences^59^) are marked by a degenerate T-rich motif, named ARS consensus sequence (*ACS*)^60, 61^ to which the heterohexameric origin recognition complex (ORC) binds^62–64^. During G1, ORC recruits the Mcm2–7 helicase to initiation sites (reviewed in Bell and Kaguni^65^), leading to the formation of the pre-replicative complex (pre-RC) that marks origin activation in S-phase (reviewed in Remus and Diffley^66^). Notably, among the >12000 high-quality *ACS* motifs, less than 300 of these function as origins of replication^67^, and only ∼120 appear to fire early in S-phase, independent of the checkpoint activation induced by dNTPs pool depletion^68^. Although the precise mechanism underlying origin selection and their single-cell temporal heterogeneity (deterministic vs. probabilistic) remains a matter of debate, their activation is thought to be modulated locally - by epigenetic modifications of the chromatin (i.e., nucleosome positioning can restrict access to the *ACS*^69^ and inhibit pre-RC assembly^70^) - and spatially in the context of the chromosome (i.e., proximity to a functional centromere^71, 72^).

Here we used a well described method to map early firing ARS regions genome-wide in yeast cell populations^68, 72, 73^. Three independent isolates of each *Hs* and *Sc* strain were synchronized in G1 using a-Factor and released synchronously in S-phase in the presence of hydroxyurea (HU), that blocks DNA elongation and causes an early S-phase arrest through dNTP starvation (Figure S4A). Prior to genome-wide sequencing, the quality of G1 and S synchronizations were evaluated by measuring DNA content using flow cytometry (Figure S4B). Mapping of the early firing ARSs was done by computing chromosome sequence coverages in early-S normalized to G1 (unreplicated control) and plotted along the reference genome at 1 kb resolution. We observed that the prominent signal corresponding to early-firing origins (indicated by black arrowheads in Figure 4A) was severely compromised and often entirely lost in *Hs* (orange plot) compared to *Sc* (blue plot). This defective firing trend was particularly obvious on the longer chromosome arms (Figure S5). Note that the reduced firing intensity of the early-S regions in the *Hs* isolates is unlikely a result of incomplete synchronization, as we accounted for their extended cell cycle and corrected with accordingly prolonged incubations (Figure S4A). The firing defect was observed genome-wide (Figure S5 and Figure 4B, ratio of origin timing in *Hs* vs. *Sc*), independent of chromosome size and ARS location (i.e., distance from the early replicating centromere). Previous high-throughput nucleosome-positioning assays have shown that well-positioned nucleosomes flanking ARS consensus sequences are conserved functional features of replication origins^29, 74^ and are maintained by ORC binding^28^. We observed that the positioning of the human nucleosomes forms the typical nucleosome-depleted region (NDR, centered on the ARS consensus); however, it is accompanied by unexpectedly higher nucleosome occupancy in the NDR-adjacent regions in both histone-humanized lineages (Figure 4C, MNase-seq profiles showing nucleosome profiles at ARSs). These results suggest that the innate increased stability of human nucleosomes^52, 53^ in yeast may have a powerful repressive effect that impinges not only the transcriptional program (shown by Truong and Boeke^50^), but also on origin firing during DNA replication. Collectively, these findings provide a mechanism to explain the previously reported cell cycle defect in the histone-humanized strains, imputed to a slow S-phase progression.

**Figure 4.**
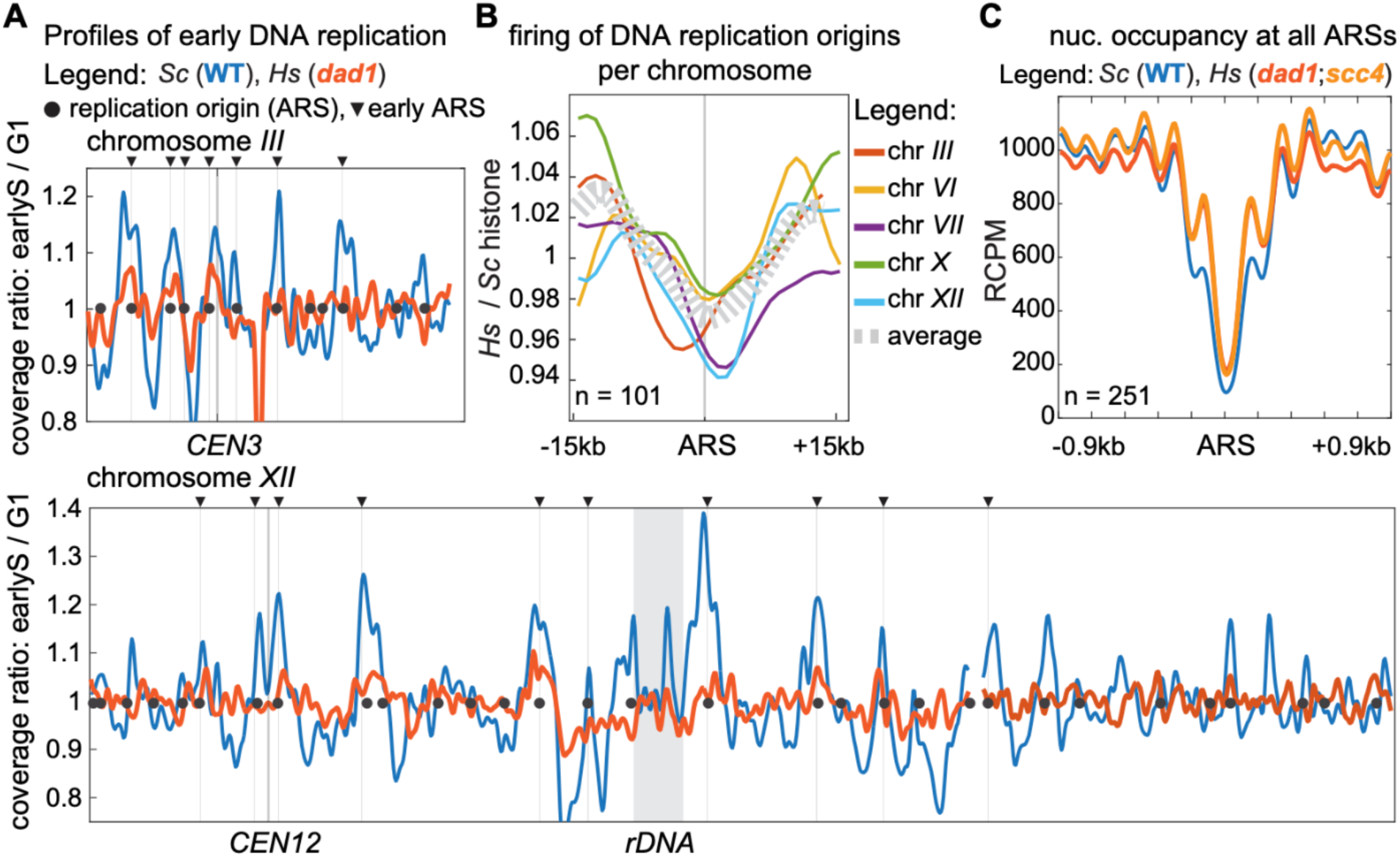
Lack of temporal activation of replication origins on humanized chromosomes. (**A**) Each track in the replication timing plots is the average representation of three independent replicates and shows the sequencing coverage ratio of early-S (HU arrested) synchronized cells normalized to the G1 (a-factor arrested) non-replicating cells (1 kb-bin size) (see also Figure S4). Replication timing profiles of the wild-type (*Sc*) are shown in blue, while those of the histone-humanized strain (*Hs,* yDT180 *dad1*-E50D) are in orange. Representative profiles of chromosome *III* (top left) and chromosome *XII* (bottom left) are shown; positions of all origins (ARS) are indicated with black circles and arrowheads indicate the early ARS subset. (**B**) Metaplots of ARS activation were computed on chromosome-by-chromosome ratios between *Hs* and *Sc* profiles (see also Figure S5) and plotted in 30 kb ARS-centered windows. (**C**) Metaplots showing nucleosome occupancy from MNase-sequencing profiles at ARSs in *Hs* (yDT180 *dad1*-E50D and yDT92 *scc4*-D65Y) compared to *Sc* strains.

### Histone humanization causes instability of the ribosomal DNA array

Intriguingly, while analyzing the deep-sequencing data (Hi-C libraries and profiles of replication timing, above), we observed a substantial enrichment in multi-mapping reads in the histone-humanized yeast (from ∼15% in *Sc* to ∼35% in *Hs* strains). Closer examination of the multimapped reads, using the built-in commands in SAMtools^75^ to sort and index the alignments, revealed that the vast majority of these originated from chromosome *XII*. In *S. cerevisiae*, chromosome *XII* harbors the highly repeated ribosomal DNA locus (rDNA; ∼150-200 copies of rRNA genes)^76^, accounting for ∼10-17% (∼1.5 Mb) of the entire yeast genome (reviewed in Kobayashi and Sasaki^77^). Given its repetitive nature and the high demand for ribosomal RNA transcripts^78^, the rDNA locus is arguably the most unstable genomic structure (reviewed in Salim and Gerton^79^). Recombination events between rDNA repeats can lead not only to variability in the size of the locus (loci)^80^ (reviewed in Kobayashi^81^), but also to the formation of extra-chromosomal rDNA circles (ERCs) thought to occur predominantly during replicative aging^82, 83^. We therefore hypothesized that histone humanization may lead to rDNA instability and copy number amplification of the rRNA genes. To test whether rDNA amplification is extra- or intra-chromosomal, we performed a Pulsed-Field Gel Electrophoresis-Southern blot assay and found an extraordinary increase to the size of chromosome *XII* (expected size ∼2.5 Mb in *Sc*) linked to the internal expansion of the rDNA locus (Figure 5A, BamHI digested chromosomes used to exclusively resolve the rDNA locus). The size of the rDNA expansion in the euploid *Hs dad1*-E50D clones exceeds the maximum resolution potential of the PFGE (5-6 Mb) but forms a band, whereas, in the aneuploid *Hs scc4*-D65Y lineage the clones display a smaller smear-like migration of the rDNA that is likely a reflection of a population of rDNAs of different sizes on the aneuploid chromosome *XII* (Figure S6A). Intra-chromosomal expansion of the rDNA is also supported by the Hi-C maps (Figure S6B, insets of ratio maps between *Sc* and *Hs* genomes showing *cis* and *trans* contact variations between chromosomes *XII* and *XIII*) and the corresponding 3D average representations of chromosomes (Figure 5B), in which the expansion of the locus causes the distal part of chromosome *XIIR* arm to be insulated from the remaining genome. Finally, we did not detect accumulation of ERCs in any of the *Hs* yeast strains compared to *Sc* (Figure S7, exonuclease treatment shows only the band of 2-micron plasmid), reinforcing the evidence of intra-chromosomal amplification of the rDNA array. To better understand the kinetics of the rDNA expansion following nucleosome humanization, we estimated the size of the locus in the euploid *Hs dad1*-E50D by computing the ratio between reads mapped to rDNA and the remainder of chromosome *XII* (“n” = number of independent genome-wide sequencing datasets, Table S2). We found that the expansion occurs very early on during histone humanization (“non-evolved” indicates genomic libraries prepared immediately after transforming the *Hs* histone plasmid and shuffling-out the *Sc* histones) and it reaches a maximum of 5-6 Mb (accounting for ∼600 repeats) after passaging them for ∼100 generations (Figure 5C, *Hs* plots). Moreover, after “re-yeastification” – by re-introducing the *Sc* histones in the already histone-humanized strains – the physiological size of the rDNA locus was entirely rescued (Figure 5C*, Hs* + *Sc* plots). Therefore, we concluded that the expansion of the rDNA locus is a reversible adaptation that is entirely dependent on human histones. Next, we wanted to learn what epigenetic-dependent mechanism(s) allows for this switch in rDNA stability.

**Figure 5.**
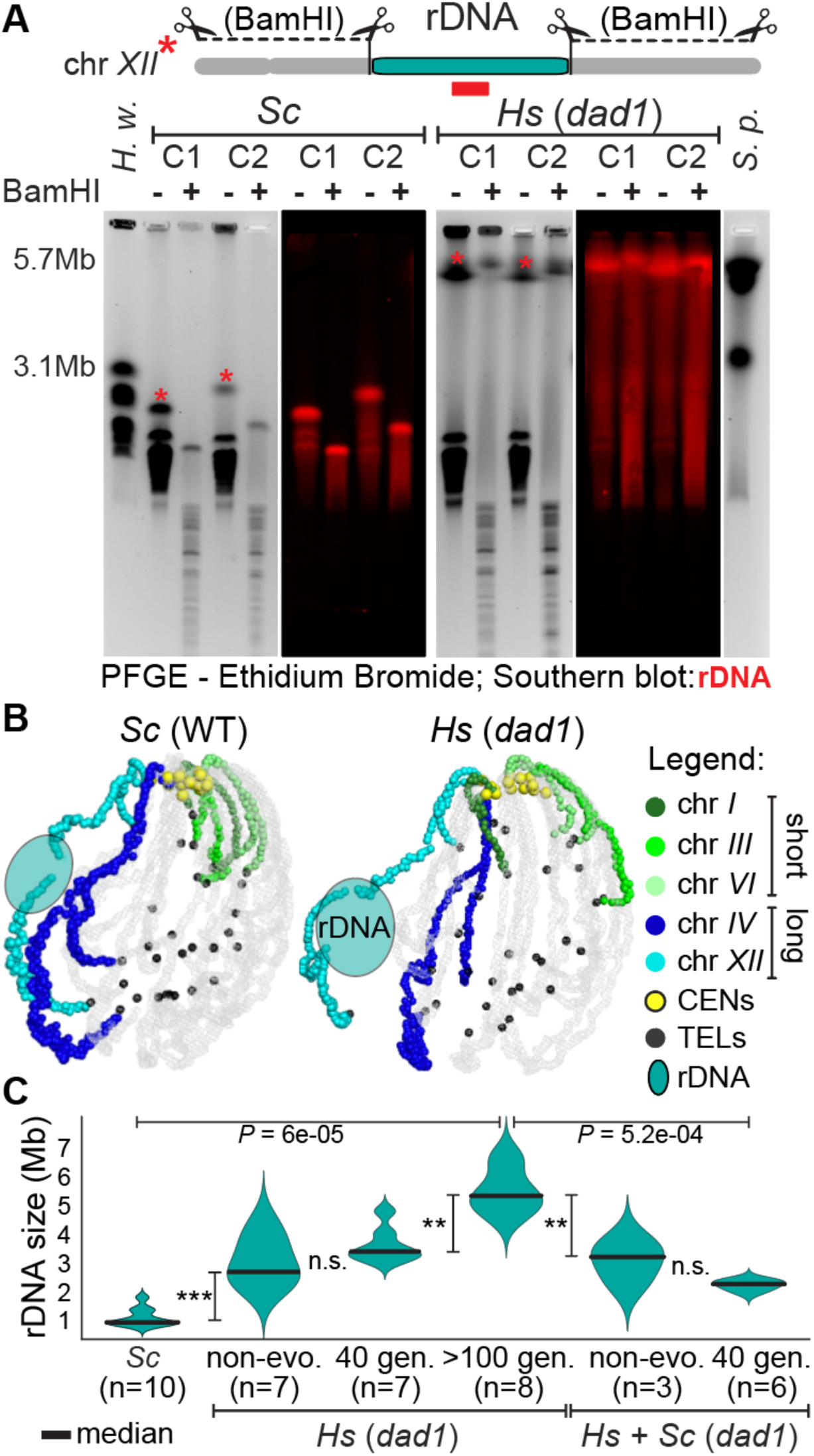
Histone humanization leads to the intra-chromosomal expansion of the repeated rDNA array. (**A**) Estimate rDNA locus sizes (turquoise region on chromosome *XII*) in *Sc* and *Hs* (yDT180 *dad1*-E50D) strains. PFGE of yeast chromosomes digested (+) or not (–) with BamHI and the corresponding Southern blot with an rDNA specific probe (red). Each “C#” represents an independent isolated clone of either *Sc* or *Hs* strain (see also Figure S6A). Left ladder: *H. wingei* chromosomes. Right ladder: *S. pombe* chromosomes. (*) indicates chromosome *XII*. PFGE run specifications: *S. pombe* program for multi-megabase chromosome separation. (**B**) 3D average representations of the *Sc* and *Hs* Hi-C contact maps (as described in Figure 3B) where the estimated position of the rDNA locus is indicated (see also Figure S6B). Color code highlight a few short and long chromosomes, as well as centromeres (CENs) and telomeres (TELs). (**C**) Violin plots showing the estimated rDNA size (Mb) calculated using rDNA-mapped reads (n = # genome sequencing datasets) in *Sc*, histone-humanized (*Hs*: “non-evo.” = non-evolved/passaged isolates; “40 gen.” and “>100 gen.” = passaged for # generations) and “re-yeastified” (native *Sc* histones added back to the humanized yeast) strains (see also Figure S9F, Table S2). *P* values were calculated using the K–S (Kolmogorov–Smirnov) test.

### Histone humanization causes rRNA metabolic dysregulation and disrupts nucleolar structure

Each ribosomal DNA repeat unit (9.1 kb) not only encodes for the four ribosomal RNA genes (*25S*, *18S*, *5.8S* and 5S), but also contains two non-transcribed intergenic spacers (*NTS1* and *NTS2*) (reviewed in Nomura et al.^84^) thought to be involved in the metabolic regulation of rDNA array size^85–87^ (reviewed in Kobayashi^81^) (Figure 6A). The amplification of this locus relies on a repeat-mediated homologous recombination mechanism that requires: (1) binding of Fob1 protein to the rDNA replication fork block (*RFB*) site^85, 86, 88, 89^ and/or (2) lack of transcriptional silencing (mediated by SIR and cohesin complexes) of the NTS sequences^87, 90, 91^. We reasoned that changes in chromatin occupancy at the rDNA locus in the *Hs* histone strains could hint towards a potential mechanism responsible for the amplification of the array. MNase-sequencing profiles showed that the *Hs* nucleosome occupancies at the rDNA locus remained unexpectedly similar between the *RDN37* and the NTS regions compared to the *Sc* yeast (where NTS silencing allows for higher nucleosome occupancy) (Figure S8A), suggesting functional misregulation. Notably we detected increased occupancy at the ribosomal origin of replication (*rARS*) in *NTS2* and at the RFB-Fob1 site in *NTS1* in the *Hs* histone strains vs. the *Sc* (Figure S8A). To validate whether RFB-Fob1 is responsible for the locus instability^86^ in *Hs* histone yeasts, we deleted *FOB1* in *Sc* strain and found that after histone humanization, rDNA arrays invariably expanded (Figure S8B). We thus conclude that rDNA amplification in the histone-humanized yeasts does not rely on a replication-based mechanism, in agreement with the absence of ERC (as previous shown in Figure S7).

**Figure 6.**
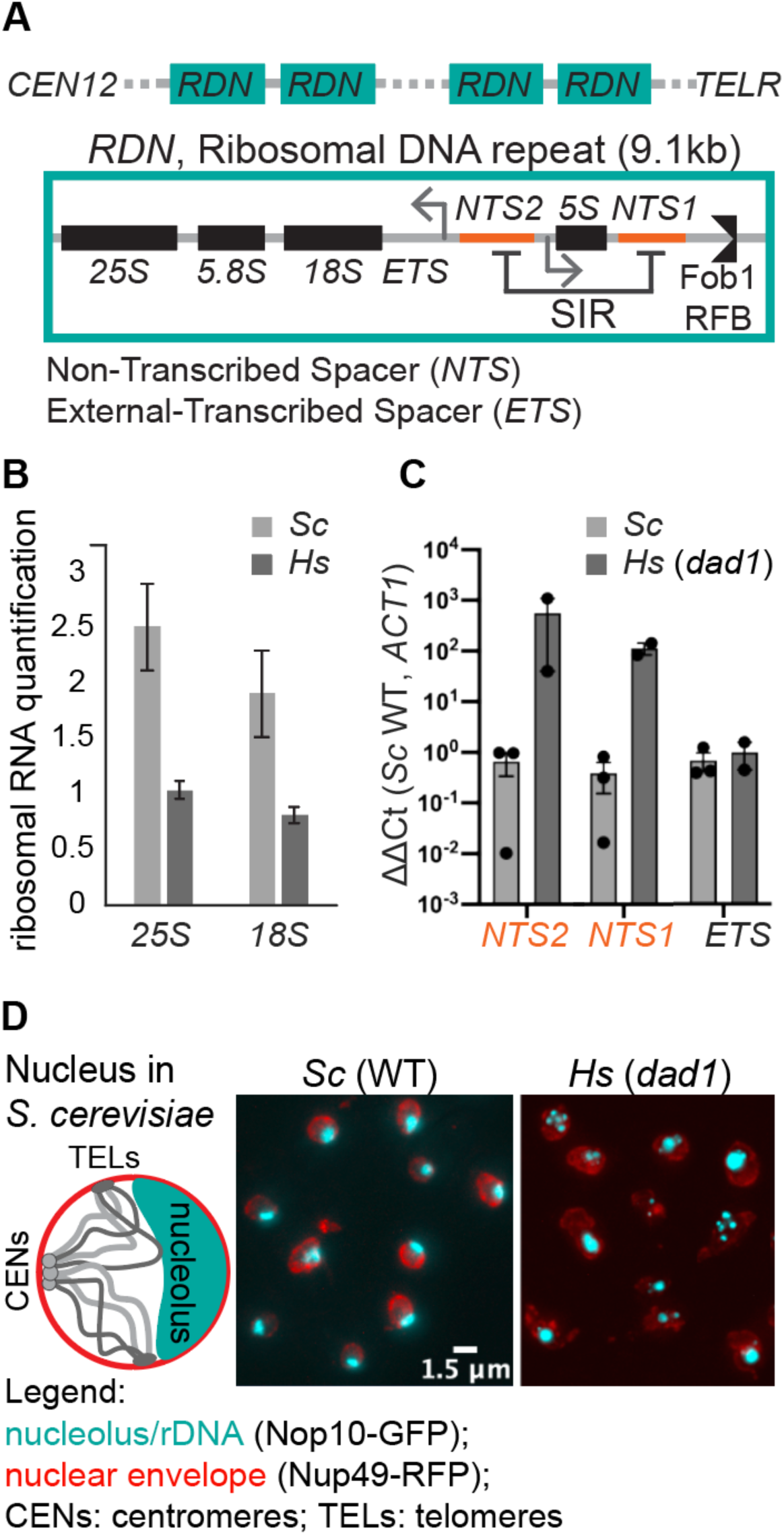
Histone humanization disrupts rDNA silencing and nucleolar structure. (**A**) Schematic showing the organization of the *RDN1* array (rDNA locus on chromosome *XII*) and an inset on an example repeat (∼9.1 kb-long), showing rRNA genes and regulatory sequences (*NTS1* and *NTS2* silenced by SIR complex, and Fob1 binding to the Replication Fork Block, *RFB*). (**B**) Quantification of rRNA levels (*18S* and *25S*) in triplicates of *Sc* and *Hs* (yDT180 *dad1*-E50D) strains. Total RNA was extracted from equivalent amounts of cells then quantified on agarose gel using ImageJ (see Figure S9B). (**C**) RT-qPCR bar plot used to estimate changes in the transcription of the *NTS1/2* and the rRNA precursor (*ETS*) relative to the housekeeping mRNA, *ACT1* (see Figure S9C). (**D**) Left, a simplified representation of nuclear organization in yeast, where examples of chromosome arms (gray lines) are anchored at the nuclear membrane through CENs and TELs, and the crescent-shaped nucleolus (turquoise) are shown. Right, representative microscopy images of *Sc* (strain: yLS110) or *Hs* (strain: yLS117) yeast nuclei. Nuclear envelope is shown in red (Nup49-RFP) and the nucleolus in cyan (Nop10-GFP).

Notably, the MNase profiles in *Hs* histone strains showed a region of nucleosome depletion mapping to the rDNA bidirectional noncoding RNA polymerase II promoter (E-pro) (Figure S8A), we thus wondered whether rRNA and/or ncRNA (at the NTSs) transcripts are dysregulated at this locus. Triplicates of total RNA extractions from similar number of cells followed by gel quantifications showed that the rRNA polymerase I transcripts (*25S* and *18S*) were ∼2.5-fold reduced in the *Hs* strains (Figure 6B and S9B). This result suggests that histone-humanized cells contain less ribosomes, in line with their substantially reduced rRNA levels. In addition to the rRNA levels, we further investigated the transcriptional activity at the E-pro by measuring the levels of NTS transcripts using RT-qPCR and RNA-seq. We detected an unprecedented increase in ncRNA at both *NTS1* and *NTS2*, observing ∼100-fold to ∼1000-fold higher levels of expression in the *Hs* strains compared to *Sc* (Figure 6C and S9C-D). The relative amounts of *ETS* (external-transcribed spacer, part of the rRNA precursor) transcripts, internally normalized to *ACT1* mRNA, remained constant between the *Hs* and the *Sc* strains, reflecting a correlation between rRNA and mRNA levels. As Sir2-dependent transcription at the E-pro has been shown to regulate rDNA copy number variation^87^, we hypothesize that the human histones in yeast are responsible for silencing defects in the NTS regions, leading to rDNA instability and locus amplification. Notably, we found that in the “re-yeastified” strains, the size of the rDNA array was reduced, and the levels of *18S* and *25S* rRNAs rebounded to their initial physiological states (Figure S9E-G).

In wild-type yeast the entire rDNA array assembles into a single subnuclear compartment, the nucleolus, forming a crescent shape structure apposed at the nuclear envelope^92^ (Figure 6D, left schematic). Previous studies have found that nucleolar localization and morphology is affected by the type of RNA polymerase (I or II) used for the rRNA synthesis^93, 94^. Moreover, nucleolar fragmentation was observed in aged yeast cells^95^, in which elevated ncRNA pol II dependent transcription^87, 91^ and variation of copy number at the rDNA locus^83^ were also detected. We thus examined whether NTS de-silencing in the histone-humanized yeasts could correlate with changes in the organization of the nucleolus. Fluorescent imaging of nuclei, using a Nop10-GFP nucleolar marker, displayed fragmentation of the nucleolus in ∼70% of the *Hs* cells compared to *Sc* (Figure 6D; see Methods for quantification). These data are consistent with predictions based on previous studies and support the role of rDNA silencing in maintain the structure of the nucleolus.

## Discussion

Our electron microscopy data suggest that the size of the *Hs-*NCP in yeast is enlarged. Since the mass of the protein/histone component of the human nucleosome is actually 0.54% “smaller*”* than that of the yeast nucleosome (109.6 kDa vs. 110.2 kDa, respectively), we conclude that the observed surface increase of the *Hs-*NCP must be due to additional nucleosome-associated DNA (corresponding to an increase of ∼10 bp in the length of the DNA protected by the NCP, see also MNase-based experiments in Haase et al., co-submission).

Given that both the nucleosome positioning and NRL remain invariant^50^, our current hypothesis is that the *Hs-*NCP is able to protect ∼10 bp more of the yeast linker DNA from MNase activity. These results support, in vivo, the model where predicted stronger interactions between the two human H2A-H2B dimers relative to yeast favor the binding stability of the human histone octamer on DNA, underlying fundamental differences in nucleosome packaging between the two species^52, 53, 55^.

Moreover, the Hi-C chromosomal maps showed that the structural effects of histone humanization go beyond single nucleosome fibers, and suggested that human nucleosomes allow for more compacted chromatin fibers in yeast (mid-range distances: 20-50 kb). These observations support a model in which higher nucleosome occupancy, accompanied by lower DNA accessibility form the basis for the drop in both mRNA and rRNA transcription and to the impairment of replication origin firing in histone-humanized yeasts. Conversely, lower humanized-nucleosome occupancy at the tRNA genes was previously shown to increase their expression^50^.

Another intriguing finding is related to the centromeric chromatin. Although the centromeric histone was not humanized (based on our investigations to date, Cse4, the yeast CenH3 specialized histone, remains unreplaceable by human CENP-A), the peri-centromeric regions (∼50 kb) appeared weakly clustered, suggesting reduced centromeric function - defined by their ability to stably segregate chromosomes. Moreover, Haase et al.^51^ showed that a specific subset of 8 centromeres (*CEN1-3*, *CEN5*, *CEN9*, *CEN11*, *CEN16*) is more frequently associated with aneuploidy, and here we found that the same set of peri-centromeres displays a <70% clustering efficiency. Therefore, it is not surprising that histone-humanized yeast lineages often display chromosomal aneuploidies (see also Haase et al., co-submission).

### Defect in DNA replication timing

The absence of a strong early S-phase origin firing in histone-humanized yeast strains resulted in a noisy temporal replication profile characterized by a multitude of small peaks, reminiscent of the stochastic/probabilistic model of DNA replication, typical of many eukaryotes, including humans^96^ and some other yeast species^97, 98^. The latter model predicts that each origin of replication follows a unique temporal program that varies stochastically from cell to cell, contrary to the deterministic one where origins have pre-established timing and frequency of firing^67, 73^. In *S. cerevisiae*, the two models can be reconciled when averaging the heterogeneous replication kinetics in a large number of cells. This has led to the postulate that the control of replication timing is deterministic at the level of large chromosomal regions but probabilistic at the level of single origins^99^. The local chromatin environment at origins was shown to affect origin activity such as both the introduction of a nucleosome within an ARS^69^ and the increased distance between nucleosomes surrounding origins^70^ led to a reduced firing. A more recent study that performed high-resolution histone chromatin immunoprecipitation followed by deep sequencing in hydroxyurea (HU) treated cells has found an inverse correlation between nucleosome occupancy surrounding origins and their firing time, that was dependent on pre-RC formation^100^. This implies that early origins with a higher frequency of ORC binding display lower nucleosome occupancy in their surroundings. These observations are particularly relevant to our work, as we detected an increase in human nucleosome occupancy in the vicinity of the ORC-binding replication origins (cumulative MNase profile at ARSs, Figure 4C), accompanied with a global loss of early firing. These results suggest that the *Hs* nucleosomes may interfere with the assembly and/or the stability of the pre-RC, which compromises the efficiency of origin firing (in agreement with previous publications ^28, 100, 101^). On the other hand, we confirmed that DNA sequences at replication origins are inherently nucleosome-disfavoring^29^, and demonstrated that this chromatin feature is independent of the NCP’s species-specificity, as it was reproduced ectopically among distantly related eukaryotes (i.e., yeast and human).

Intriguingly, humanized Orc4 subunit was shown to cause the loss of ORC’s selectivity for ARSs, leading to its promiscuous and stochastic binding to the constitutively open chromatin of yeast^102^. Our work showed that the less accessible histone-humanized yeast chromatin loses its characteristic deterministic replication program. Therefore, we cannot exclude the possibility that the replication defect in the histone-humanized yeasts maybe due to the increased stability of the *Hs-*NCP that may impinge on the activity of nucleosome/chromatin remodelers. Alternatively, in light of the remarkable instability of the rDNA locus in the humanized yeasts, DNA replication of the expanded rDNA locus may require the recruitment of an excess of limiting replication initiation factors^103^, causing their widespread depletion at replication origins throughout the rest of the genome.

### Expansion of the ribosomal DNA locus

Notably, about half of the rDNA repeats are transcriptionally active at any one time^104, 105^. Work from Ide et al.^106^ in budding yeast, established the importance of the extra, untranscribed rDNA repeats as “protective” against DNA damage in the highly transcribed array. The authors concluded that while the extra copies of rDNA may not be essential to meet cellular rRNA demands, rather, they may serve to reduce the transcriptional load on the rDNA to allow replication-coupled repair and maintain the integrity of this essential locus, especially under stress conditions. In our case, histone humanization may be seen as a source of endogenous stress leading to drastic transcriptional dysregulation of the NTS sequences, accompanied by a remarkable increase in rDNA gene copy number (Figure 5-6). rDNA expansion appeared entirely intra-chromosomal (as we failed to detect enrichment in extra-chromosomal rDNA circles, ERC, Figure S7), it is thus plausible that rDNA array expansion and concomitant reduction in rRNA transcripts (RNA polymerase I) represent genomic adjustments necessary to counterbalance exacerbated transcriptional activity caused by lack of silencing at the NTS (RNA polymerase II). In other words, the rDNA expansion may serve as a reservoir of RNA pol I inactive genes to release the overall transcriptional burden and maintain the integrity of this essential locus (in agreement with Ide et al.^106^). This hypothesis is supported (1) by the rapid and consistent rDNA size adjustments, when histone genes are swapped from *Sc* to *Hs* and vice versa Table S2B-C, and (2) by higher nucleosome occupancy at the *rARS*, which may lower its firing efficiency thus reducing transcription-replication fork collision that leads to DNA damage response, recombination and ERC formation.

We have shown that the expansion of the rDNA locus is due to silencing defects. However, it remains unclear how human histones interfere with this process given that the Sir2 deacetylated lysine residues on the H3 and H4 are conserved and that none of the SIR factors (silencing: Sir2, Sir3, Sir3) nor the RNA I/II/III pol genes were found to be differentially expressed in *Hs* vs. *Sc* (Table S4). As often times chromatin modifying enzymes (e.g., SIR factors) require to contact extensive patches on the surface of the nucleosomes^107^, we cannot exclude that cumulative changes introduced by the *Hs*-NCP may disrupt these interactions and affect their downstream functions in yeast.

### A potential interplay between cell size and transcriptional changes

Biosynthesis of total RNA and proteins increases in proportion to cell size such that their concentrations remain approximately constant as a cell grows (reviewed in Xie et al.^108^). This size-dependent transcriptional scaling is thought to ensure constant concentrations of total mRNA, rRNA and tRNA, to regulate protein synthesis in proportion to cell size (reviewed in Marguerat and Bahler^109^). An intriguing model for rDNA copy number regulation has proposed that Sir2 activity (the NTS silencing factor implicated in the stability of the rDNA array) may decrease after cell enlargement, allowing the increased recombination at the rDNA and its expansion^110^. Furthermore, recent works by Swaffer et al.^111^ and Sun et al.^112^ showed that the increase of RNA polymerase II initiation rate is the major limiting factor for increasing transcription with cell size in yeasts. Here we hypothesize that the cell size increase observed in the histone-humanized yeasts^50^ may be correlated with the extraordinarily high transcriptional activity of the RNA pol II at the E-promoter in the NTS regions of the rDNA (Figure S9C-D). Our current model predicts that the lack of silencing at the NTS will titrate more RNA pol II, causing its depletion from the free inactive pool whose feedback may eventually translate into both a cell size increase and rDNA expansion. Experiments to assess both the occupancy of the RNA pol II and the molecular crowding in the histone-humanized yeast cells are required to validate this model. We expect to observe that the increased RNA pol II occupancy at the rDNA is anticorrelated with molecular crowding, given that transcriptional excess does not lead to functional mRNAs nor rRNA involved in translation.

Finally, several studies have found that chromatin remodelers (SMC), such as cohesins and condensins bind many locations in the yeast genome^113, 114^, where they play important roles in the organization of the chromatin. Relevant examples are the origins of replication^115^, the pericentromeric regions and the nucleolus (reviewed in Lawrimore and Bloom^116^), where SMCs are involved in preserving rDNA stability by presumably maintaining silencing at the *cis*-intergenic sequences^87, 91^. As, our results showed an unprecedent increase in non-coding RNA transcription at the *NTS2*, we cannot exclude the possibility that the human nucleosome in yeast may affect centromere clustering, firing of replication origins and rDNA stability by altering the higher-order SMC-dependent organization of the chromatin.

## Acknowledgments

We thank Sarah French (UVA) for extensive guidance on chromatin preparation for TEM and the microscopy facility at NYU Langone for electron microscopy training. We thank David Truong (NYU Tandon), Ran Brosh (NYU Langone), Julien Mozziconacci (MNHN), Jeffrey Smith (UVA) for helpful discussions and comments on the manuscript, and the entire Boeke lab for their assistance. This work was supported in part by National Science Foundation grant MCB-1921641 to J.D.B.

## Author Contributions

L.L.-S. and J.D.B. designed the research; L.L.-S. and M.A.B.H. performed the experiments and analyzed the data; L.L.-S., M.A.B.H. and J.D.B. wrote the manuscript.

## Methods

### Resource availability

#### Lead Contact

Further information and requests for resources should be directed to Jef D. Boeke.

#### Materials availability

Yeast strains generated in this study can be requested directly by contacting the lead contact. This study did not generate new unique reagents.

#### Data and code availability

Raw microscopy images were deposited on Mendeley DOI: 10.17632/2j5pzfm2xm.1 FASTQ files of GWS (HiC datasets and RNA sequencing) in were deposited in the NCBI GEO database.

BioProject: PRJNA951416

https://urldefense.com/v3/ https://dataview.ncbi.nlm.nih.gov/object/PRJNA951416?reviewer=9gl1a491djdustacitl4trn5qr;!!MXfaZl3l!fXrF-4m-0vTACLrafzreljsCgI5nK8nx-v4DByrn13QNAIgES-GpMkf7kLQEo5QQ_d05EHES8O_t-QRgCu1Cz5APaqcRSf9Z$ https://dataview.ncbi.nlm.nih.gov/object/PRJNA951416?reviewer=9gl1a491djdustacitl4trn5qr

No new code was generated in this study.

### Method details

#### Experimental models and subject details

Yeast strains used in this work are listed in the resource Table S5. The deletion of the *FOB1* coding sequence was achieved using CRISPR-Cas9 in the “shuffle strain” (yMAH666 in which the encoding yeast histones are exclusively on a centromeric plasmid), and the oligonucleotide sequences used as gRNA and repair donors are provided in the resource Table S5.

#### Media and culture conditions

All strains listed were grown in rich medium (Yeast extract Peptone Dextrose (YPD): 1% bacto peptone (Difco), 1% bacto yeast extract (Difco) and 2% dextrose) liquid or solid (2% agar) at 30°C unless otherwise specified in the methodology below.

Growth curve assay after “histone re-yeastification”

Yeast cultures from three independent isolates of the *Sc* histone strain (yDT67) and 3 of the reyeastified *Hs* stains (*Sc* yMAH753/4/5 and *Hs* + *Sc* yMAH756/7/8) were grown to saturation in YPD liquid medium at 30°C. Yeast cultures in stationary phase were diluted in fresh YPD medium to an optical density (OD) A600 = 0.07, 200 µl were transferred to 96 well plates and every minute the BioTek Eon microplate spectrophotometer was programmed to shake the plate and measure the OD600 every 15 min for a total of 24 h at 30°C. OD600 values were imported in GraphPad Prism version 9 for Mac OS (GraphPad Software, San Diego, California USA, www.graphpad.com) and used to calculate mean and standard deviation for each isolate of each strain.

#### Transmission electron microscopy (TEM) for imaging chromatin fibers

For the preparation of chromatin spreads in yeast we followed the published protocol described by Osheim et al.^117^. We extracted and spread chromatin from log phase yeast cultures (*Sc* histones: BY4742; *Hs* histones: yDT92, yDT180) grown in YPD with 1M sorbitol at 30°C. Approximately 10^7^ cells were enzymatically lysed using a 1mg/ml Zymolyase 20T (US biological, Z1000) solution in YPD 1M sorbitol. Chromatin spreading was conducted in a 35 x 10 mm plastic petri dish containing a 0.025% Triton pH 9.1 solution that was incubated at room temperature and in mild agitation for 45 min. Spreading chromatin was mildly crosslinked with 1/10 [v/v] sucrose–formalin solution (100 mM sucrose, 3.7% formaldehyde Tousimis Researc Corporation, 1008A, with the pH adjusted to 8.8) for an additional ∼30 min. Chromatin was deposited onto the EM carbon grids (Electron Microscopy Sciences, CF300-Cu) by centrifugation at 7000 x g (Centrifuge: Sorvall LXTR with swinging bucket rotor) for 10 min. Nucleic acid and protein staining were performed using 4% solutions of Uranyl Acetate (UA) (Electron Microscopy Sciences cat. 22400-4) and Phosphotungstic acid hydrate (PTA) (Sigma-Aldrich P4006-10G) in ethanol. Imagese were acquired using the electron transmission microscope (FEI Talos 120C TEM) at various resolutions, ranging from 10-kX to 150-kX, at the NYU Langone Microscopy Laboratory.

#### Estimating surface area of the nucleosome core particles (NCPs)

Mononucleosome size was measured using images acquired at 200 nm and 100 nm resolution using ImageJ^118^. Prior measuring of each image was calibrated on the scale bar provide in the electron microscope image. Raw values can be found in Table S1.

#### Hi-C: library preparation

Hi-C experiments and data analysis were performed as described^72, 119^ unless otherwise indicated in the following method description. Briefly, independent yeast isolates were inoculated into 5 ml YPD medium and grown overnight at 30°C. The following morning the overnight cultures were subcultured into 150 ml fresh YPD for ∼3 h at 30°C until reaching ∼1.2 x 10^9^ cells total (∼120 OD). Cells were crosslinked using 3% [v/v] formaldehyde for 20 min at room temperature and then quenched with 350 mM glycine for 15 min at 4°C in mild agitation. Crosslinked cells were harvested by centrifugation at 1500 x g for 5 min at 4°C, washed twice with cold fresh medium, and resuspended in 5 ml spheroplast solution (1M sorbitol, 50 mM potassium phosphate, 5 mM DTT, 250 U zymolyase 100T [US Biological, Z1004]) for 50 min incubation at 30°C. Spheroplasts were harvested by centrifugation at 2500 x g for 10 min at 4°C, washed with 10 ml of cold 1 M sorbitol and resuspended in 2 ml of 0.5% SDS, H_2_O at 65°C for 20 min. The crosslinked chromatin was enzymatically fragmented using 125 U of MboI (NEB, R0147) in a final reaction volume of 3 ml (1X Cutsmart NEBuffer, 0.33% SDS and 2% Triton) and an incubation at 37°C overnight (up to 16 h). The digested product was centrifuged at 18000 x g for 20 min and the pellet was resuspended in 200 μL cold water. DNA sticky ends were filled in (to blunt ends) using a biotin-labeled 30 μM dNTP mix (dATP, dGTP, dTTP and Biotin-14-dCTP Thermo Fischer, 19518018) and Klenow enzyme (NEB, M0210L) at 37°C for 80 min. Biotinylated restriction fragments were re-ligated using 60 Weiss Units of T4 DNA ligase (Thermo Fischer, EL0014) in 1.2 ml final volume at room temperature for 2 h in mild agitation. Ligation product was reverse cross-linked by 0.5 mg/mL proteinase K (Thermo Scientific, EO0492) in 0.5% SDS, 25 mM EDTA buffer at 65°C for 4 h. The un-crosslinked sample was ethanol precipitated and purified using the large fragment DNA recovery kit (Zymo Research, D4046). Religated-biotinylated restriction fragments were pulled down using Dynabeads MyOne Streptavidin C1 magnetic beads (Invitrogen, 65001) according to the manufacture protocol. The final cleaned-up Hi-C library was used as input material for Illumina sequencing library prep kit (NEB, E7805) with 6-8 cycles of PCR amplification using KAPA-HiFi (Kapa Biosystems, KK2602). DNA library was sequenced using an Illumina NextSeq 500 75-cycle high output kit.

#### Hi-C: data processing

To generate contact maps: paired-end reads were processed using the HICLib algorithm^120^ adapted for the *S. cerevisiae* genome. Read-pairs were independently mapped using Bowtie 2^121^ (mode: --very-sensitive --rdg 500,3 --rfg 500,3) on the corresponding reference sequence^122^ (S288c available on SGD) indexed for MboI restriction site. In the contact frequency maps, the unwanted restriction fragments (RFs) were filtered out (e.g., loops, non-digested fragments, etc.; as described by Cournac et al.^123^), whereas, the valid RFs were binned into units of fixed size bins of 5 kb. Bins with a high variance in contact frequency (<1.5 S.D. or 1.5–2 S.D.) were discarded to remove potential biases resulting from the uneven distribution of restriction sites and variation in GC% and mappability. The filtered contact maps were normalized using the sequential component normalization procedure (SCN)^123^. Approximately 10-15 million valid contacts were used to generate a genomic contact map for each triplicate.

#### Contact probability in function of the genomic distance, p(*s*)

The Hi-C contact probability (p) decreases as the genomic distance (*s*) between restriction fragments increases^119^. p(*s*) plots were computed on intra-chromosomal read pairs from which self-circularizing and uncut events were discarded^123^. The retained reads were log-binned in function of their distance along chromosome arms, such as the p(*s*) shows the distribution of the sum of contacts weighted by both bin-size 1.1^(1+bin)^ and chromosome length (*s*). Comparison of the degree of p(*s*) decay is indicative of a change in polymer state.

Log2 ratios of Hi-C contact maps are used to detect contact variation between genomes^119^. Each pairwise comparison was computed on Hi-C normalized maps binned at 5 kb and the log2-ratio map was Gaussian smoothened (window size of 50 kb).

For the 3D representations we used the “Shortest-path Reconstruction in 3D” (ShRec3d)^124^ algorithm as previously described^72^. Finally, the average genome structures were visualized using PyMol.

#### Pulsed-field gel electrophoresis (PFGE) and Southern blotting

Chromosomes from stationary yeast cultures (*Sc* histones: BY4741, BY4742, yDT67, yMAH1242-12447; *Hs* histones: yDT92, yDT180, yLS118-123) were prepared in agar molds using the Certified Megabase Agarose (Bio-Rad, 1613108), and PFGE was carried out with running conditions recommended for *S. pombe* chromosomes (BioRad, 170-3633) to maximize size resolution of the largest chromosomes, as described previously^125^. In agar chromosome digestion with BamHI (NEB, R0136L) was used to release the entire rDNA locus (∼1.5 Mb to ∼5 Mb) from chromosome *XII*. Agar molds treated or not with BamHI were then used for the PFGE and Southern blot. These methods were reported in detail in our previous publication Lazar-Stefanita et al.^80^. In this specific experiment, we used oligos mapping in the *ETS* and *18S* sequences of the *RDN37* repeat to generate by PCR a DNA probe (769-bp long), that was labelled using Klenow Fragment exo- (NEB, M0212L) with Digoxigenin-11-dUTP alkali-stable (Roche, 11093088910) at 37°C. The labeled and denatured probe was used for the Southern blot hybridization on a nylon membrane (Pall® 60208 BiodyneTM B Membrane, 60208) containing the transferred DNA from the PFGE. A primary rabbit anti-DIG antibody (working concentration 1:4000 in Blocking buffer Odyssey; ABfinity™ Rabbit Monoclonal, 700772) followed by a secondary antibody (working concentration 1:10000 in Blocking buffer Odyssey; LIRDye® 680RD Goat anti-Rabbit IgG (H + L), 0.5 mg, 926-68071) were used to specifically detect the rDNA locus using a LI-COR Odyssey® Imager.

#### Exonuclease treatment to detect Extra-chromosomal rDNA circles (ERC)

Genomic DNA was extracted from agar plugs (as described above for PFGE chromosome preparation) using the Zymoclean gel DNA recovery kit (Zymo Research, D4001T) and successively digested with Exonuclease V (RecBCD, NEB M0345) at 37°C for 3 h. Circular plasmid (pUC19, NEB N3041S) and sheared (sonicated) genomic DNA were used as digestion controls.

#### Cell cycle synchronization and DNA staining for flow cytometry

G1 arrested cells were obtained in triplicate by incubating log-phase growing *Sc* (yDT67) and *Hs* (yDT180) strains (OD600= 0.3 - 0.5; ∼10^7^ cells/ml) in YPD supplemented with 0.1 μg/ml a-factor (Zymo Research, Y1004) for 3 h 30 min (yDT67) or 4 h 30 min (yDT180) at 30°C. Aliquots of ∼2 x 10^7^ G1 cells were fixed in 70% ethanol to asses synchronization efficiency; while, the remainders were centrifuged, washed twice with fresh medium and finally resuspended in medium containing 200 mM hydroxyurea (HU; Sigma-Aldrich, H8627-25G). These latter cultures were incubated for 1 h 30 min (yDT67) or 3 h 30 min (yDt180) at 30°C and aliquots were sampled to microscopically assess for early S-phase arrest.

All G1 and HU aliquots (∼10^7^ cells/replicate, fixed in 70% ethanol) were stored at 4°C overnight and successively processed for DNA content analysis using flow cytometry. Cells were pelleted (at 3000 x g for 3 min) and washed three time with 2 ml of RNAse solution (10 mM Tris pH 8.0, 15 mM NaCl) before being treated with 0.1 mg/ml RNAse A for 3-4 h at 37°C. Cells were washed once with 50 mM Tris pH 8 and resuspended in labeling solution (1 μM SYTOX Green in 50 mM Tris pH 8; Thermo Fisher) for 1 h at 4°C protected from light. Before flow cytometry data acquisition, cells were washed three times and resuspended in 50 mM Tris pH 8. Flow cytometry was performed on a BD Accuri C6 Flow Cytometer (BD CSampler Software) and data analyzed using FlowJo v10.0.7 software.

#### DNA replication timing

Each profile of replication timing was generated from three independent clones of *Sc* (yDT67) and *Hs* (yDT180) strains (see: cell cycle synchronization and DNA staining for flow cytometry) by deep-sequencing analysis as described previously^73^. Briefly, fractions of replicating and non-replicating cells were obtained by arresting cells with a-factor for 3 h 30 min (yDT67) or 4 h 30 min (yDT180), then they were washed and released in HU for 1 h 30 min (yDT67) or 3 h 30 min (yDt180) at 30°C. Synchronization efficiencies were validated by flow cytometry. Pellets of ∼6 x 10^8^ cells were used to extract genomic DNA using acid-washed beads (Sigma-Aldrich, G8772-100G) and phenol-chloroform (Thermo Scientific). Library preparation was performed using the NEBNext Ultra II FS kit (NEB, E7805L) according to the manufacturer’s protocol. Resulting libraries were paired-end deep-sequenced (2 x 36 bp cycles) on NextSeq500 Illumina platform. Reads were mapped to the corresponding reference genome using Bowtie 2^121^ in its --very-sensitive mode. Profiles of replication timing were generated by normalizing the replicating (S-phase, HU) sample to the non-replicating (G1, a-factor) sample in 1 kb bins. The resulting ratios were Gaussian-smoothed (window size of 10 kb) and plotted by genomic coordinate, measuring variations in DNA copy number as a proxy of replication time.

#### Nucleosome maps at replication origins and ribosomal DNA locus

We used published MNase-seq datasets^50^ to evaluate nucleosome occupancy in the proximity of replication origins and at the rDNA locus. Genome-wide positions of replication origins, defined as ORC-binding sites with ARS consensus sequence (total ARS = 251), were obtained from Eaton et al.^29^. Nucleosome maps were generated following the methods described in the co-submitted work by Haase et al.

#### RNA extraction

Total RNA was extracted from 3 independent isolates of *Sc* (yDT67), *Hs* (yDT180, yDT92) and re-yeastified *Hs* (*Sc* yMAH753/4/5 and *Hs* + *Sc* yMAH756/7/8) strains. Approximately 2 x 10^8^ cells were harvested from mid-log phase cultures (1.5-2 x 10^7^ cells/ml) grown in YPD medium at 30°C. Cell pellets were washed in RNase free water and resuspended in RNA lysis buffer (50 mM Tris-HCl pH 8, 100 mM NaCl). Cells were lysed mechanically using acid-washed glass beads (Sigma-Aldrich, G8772-100G) at 4°C. The RNA was extracted by phenol:chloroform:isoamylalcohol (ThermoFisherScientific, 15593) and ethanol precipitated. Extractions were treated with DNaseI (Agilent, 600031) for 1 h at 37°C and RNA quality was verified by agarose gel in 1X TAE.

#### RNA-based assays

Reverse Transcriptase (RT) - quantitative PCR assay. Triplicates of total RNA extractions from *Sc* (yDT67) and *Hs* (yDT180) strains were used for RT-qPCR reactions with gene specific oligos (rRNA: *NTS1*, *NTS2*, *ETS1*; mRNA: *ACT1*)^91, 126^. The RT reaction was performed according to the manufacturer protocol SuperScript™ IV Reverse Transcriptase (Invitrogen, 18090050). Successively, quantitative PCR was performed using the LightCycler® 480 SYBR Green I Master (Roche, 04887352001) following the standard amplification protocol with 45 cycles in a multi-well PCR plate 384. Ct values for each replicate were imported in GraphPad Prism version 9 for Mac OS (GraphPad Software, San Diego, California USA, www.graphpad.com) and used to calculate mean and standard deviation for each gene in each strain. Raw Ct values can be found in Table S3. For RNA-seq data and analysis (Figure S9D and Table S4) refer to the co-submitted work by Haase et al.

#### Protein tagging and Fluorescent microscopy

The organization of the nucleolus within the nucleus was monitored using fluorescently tagged proteins at their endogenous C-terminus. Nuclear envelope was labeled with mScarlet (*NUP49::mScarlet-S.p. HIS5*) and the nucleolus with GFP (*NOP10::EGFP-KanMX*) using reagents that we previously described in Lazar-Stefanita et al.^80^ (see strains in the resource Table S5). Two independent isolates for each strain, containing either *Sc* or *Hs* histones, were validated for dual tagging based on their positive emission wavelengths in the GFP (513 nm) and RFP (605 nm) channels. The resulting strains (*Sc*: yLS110-C1 and yLS110-C3; *Hs*: yLS117-C1 and yLS117-C2) were grown in SC–His medium to saturation (24 h for yLS110 and 48 h for yLS117) and live cells were imaged in agarose pads prepared in SC–His medium (to prevent Brownian motion). Imaging was performed on the EVOS M7000 microscope using the Olympus X-APO 100 Oil, 1.45NA/WD 0.13mm (Oil) objective. Images were acquired as Z-stacks and visualized as max intensity projections using ImageJ^118^. Different fields of view were used to count nearly 1000 nuclei (496 for yLS110 and 477 for yLS117) displaying either one intact nucleolus or many fragmented nucleoli.

### Quantification and statistical analysis

Information on the number of biological replicates, statistical tests and *P* values are provided in the Method details and Figure legends.

## Supplemental Figure Titles and Legends

**Figure S1, related to Main Figure 1.**
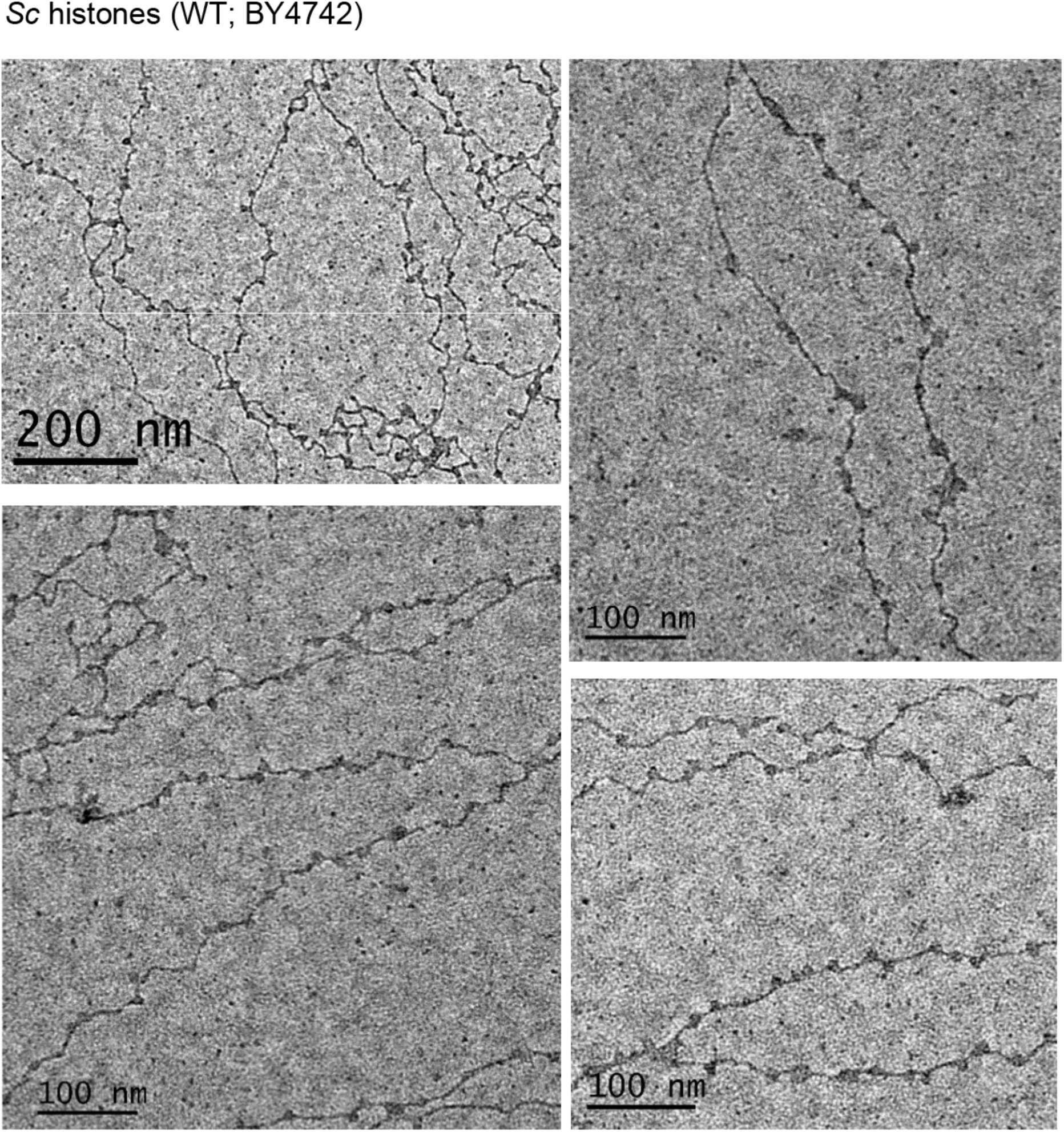
Nucleosome fibers of wild-type yeast with native histones (*Saccharomyces cerevisiae, Sc*, strain: BY4742). Representative panels showing the 10 nm fibers at different resolution (scale bars: 100 nm and 200 nm).

**Figure S2, related to Main Figure 1.**
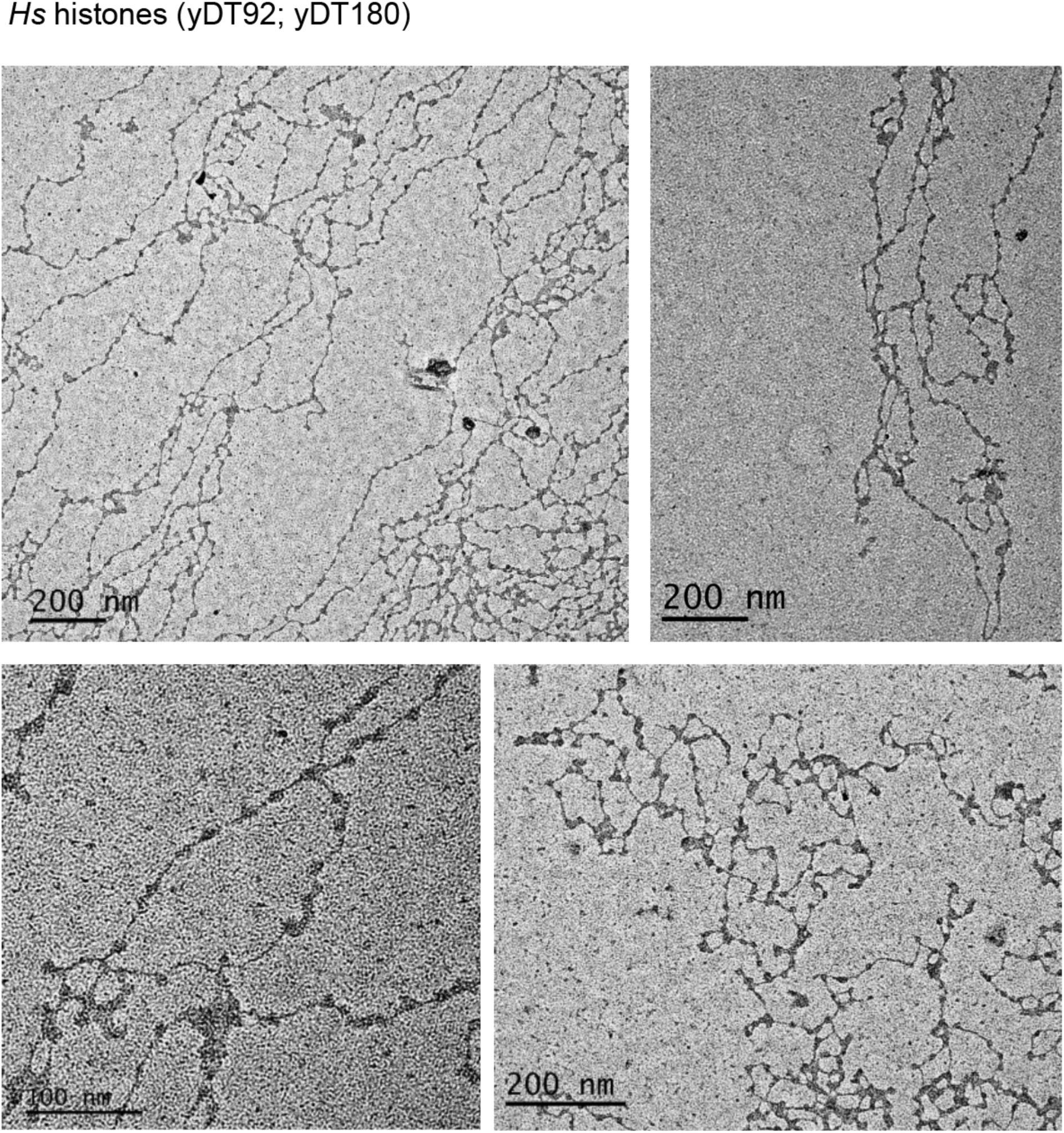
Nucleosome fibers of histone-humanized yeasts (*Homo sapiens, Hs*, strains: yDT92, yDT180). Representative panels showing the 10 nm fibers at different resolution (scale bars: 100 nm and 200 nm).

**Figure S3, related to Main Figures 2 and 3.**
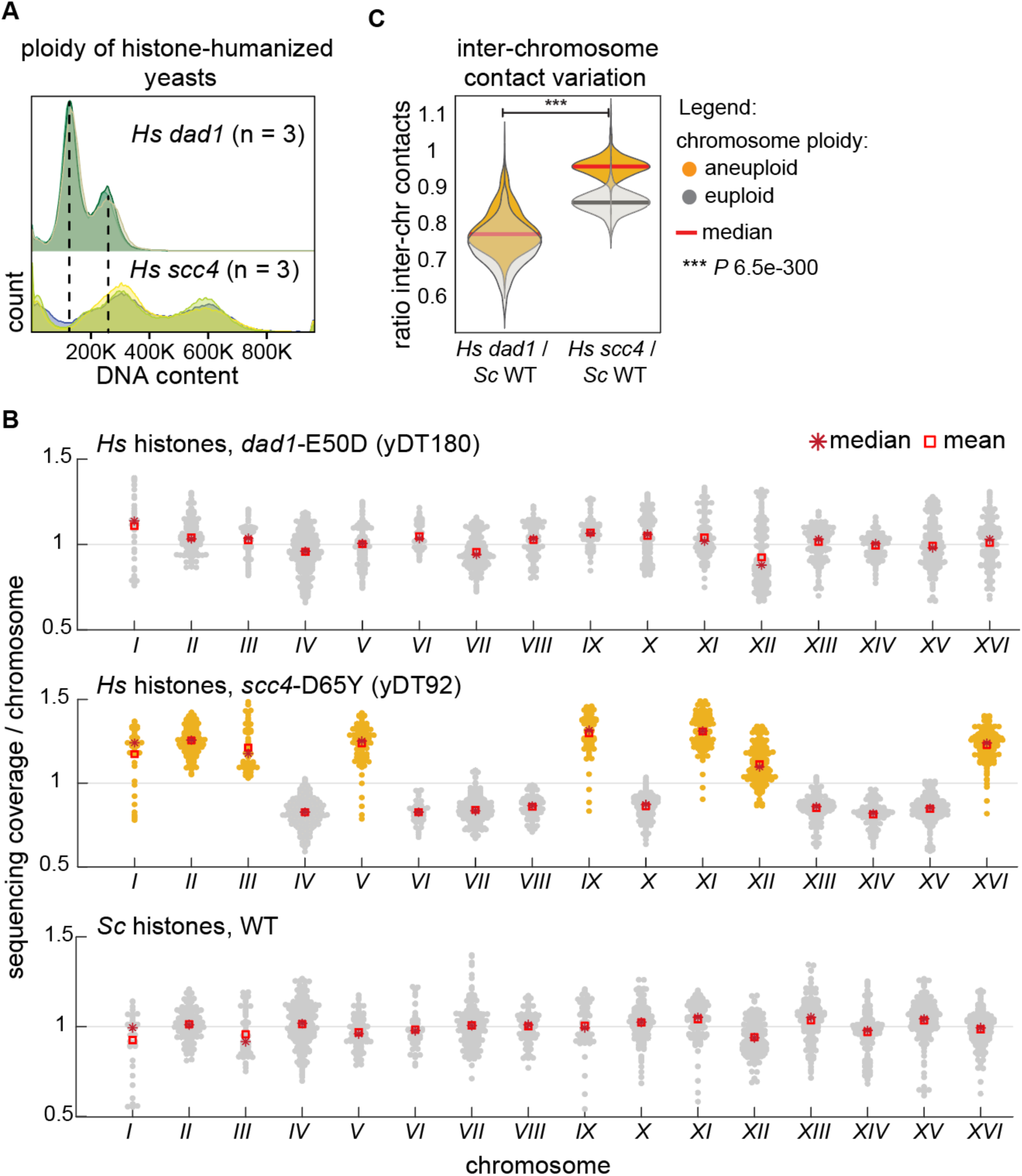
Ploidy varies among the histone-humanized strains. (**A**) Flow cytometry histograms showing DNA content in histone-humanized yeast strains stained with SYTOX Green. *Hs* euploid: yDT180 *dad1*-E50D; *Hs* aneuploid: yDT92 *scc4*-D65Y. (**B**) Average of chromosome sequencing coverage normalized by the total number of reads. Aneuploid chromosomes (increased copy number) are shaded in amber. (**C**) Inter-chromosome contact variation in the histone-humanized genomes (*Hs*) relative to wild-type (*Sc*). Normalized Hi-C contact maps (complete maps of the insets shown in Figure 2A) were used to compute the ratios between *Hs* and *Sc* strains, which were then plotted according to the level of chromosome ploidy (aneuploid vs. euploid). The increase of intra-chromosome contacts in the *Hs* strains (Figure 2B-C) is likely responsible for the ratio < 1 observed in both the euploid (yDT180) and in the non-aneuploid chromosomes of yDT92, as an effect of the normalization process. *P* values were calculated using the K–S (Kolmogorov–Smirnov) test in MATLAB 2018.

**Figure S4, related to Main Figure 4.**
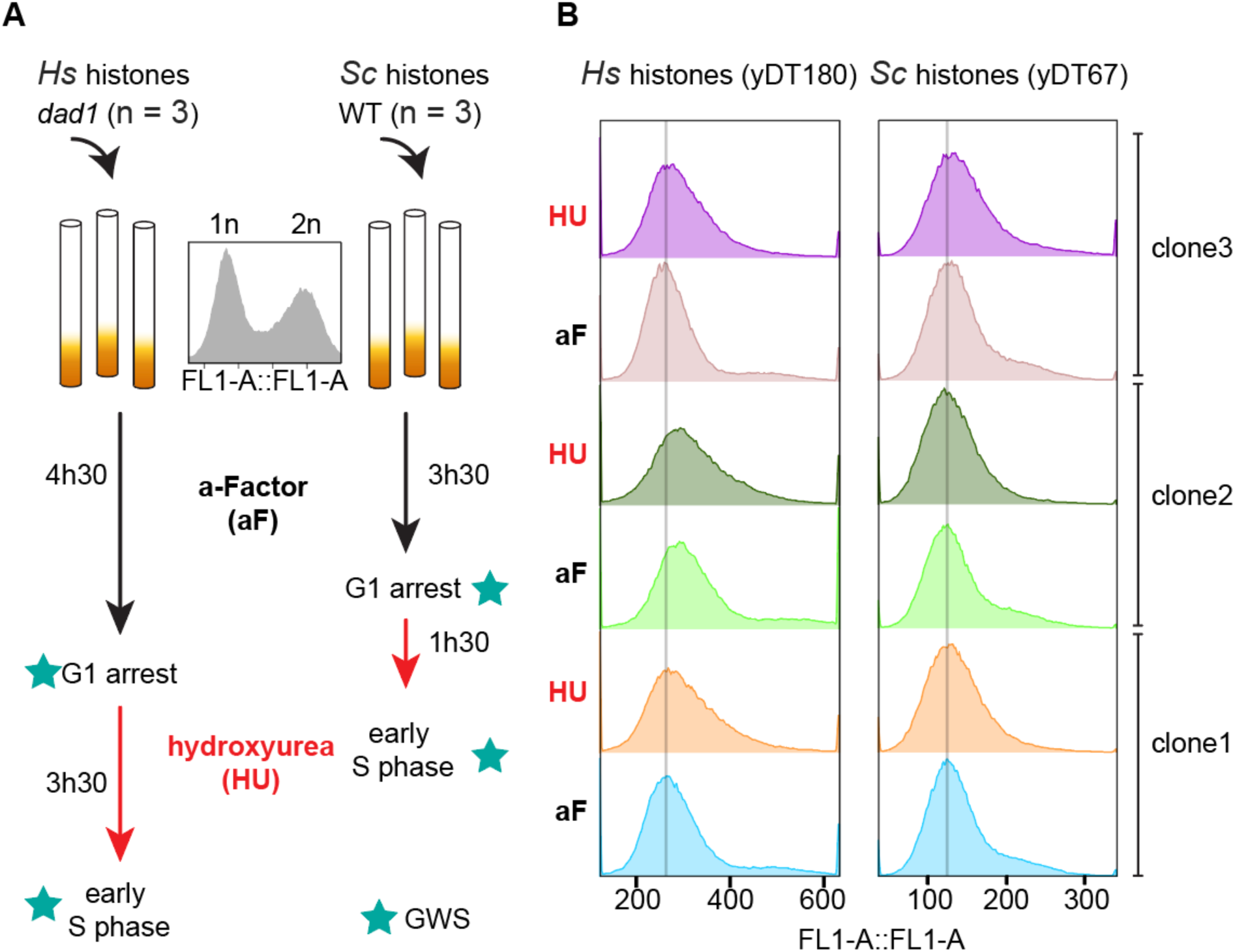
Method for mapping replication timing in wild-type and histone-humanized yeasts. (**A**) Schematics of the experimental approach used to grow and synchronize yeast cells with either native (*Sc*) or human (*Hs*, strain: yDT180 *dad1*-E50D) histones in G1 and early S phase. Star-labeled steps indicate genome-wide sequenced samples used to generate replication timing profiles. (**B**) Flow cytometry histograms measuring DNA content of the three independently synchronized cell cultures in A, stained with SYTOX Green. As expected, no obvious differences are observed between G1 and early-S phase synchronized cells.

**Figure S5, related to Main Figure 4.**
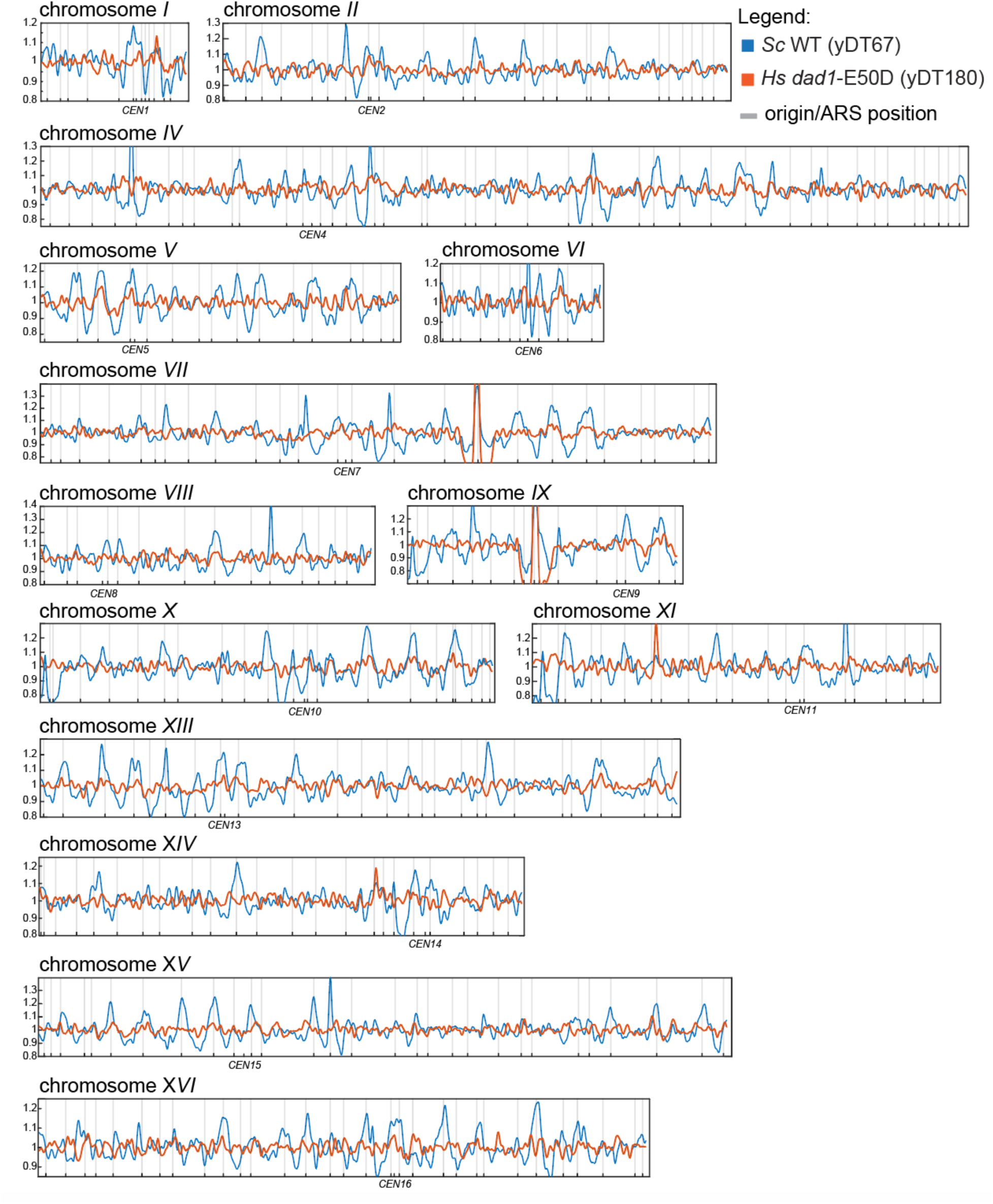
Genome-wide replication timing profiles in wild-type and histone-humanized yeast strains. Each track in the replication timing plots is the average representation of three independent replicates and shows the sequencing coverage ratio of early-S (HU) synchronized cells normalized on the G1 (a-factor) non-replicating cells (1 kb-bin size). Chromosome-by-chromosome replication timing profiles of the wild-type (*Sc*) are shown in blue, while those of histone-humanized (*Hs,* yDT180 *dad1*-E50D) are in orange. Origin (ARS) positions are indicated with gray vertical lines and centromere (CEN) positions are indicated below each plot.

**Figure S6, related to Main Figure 5.**
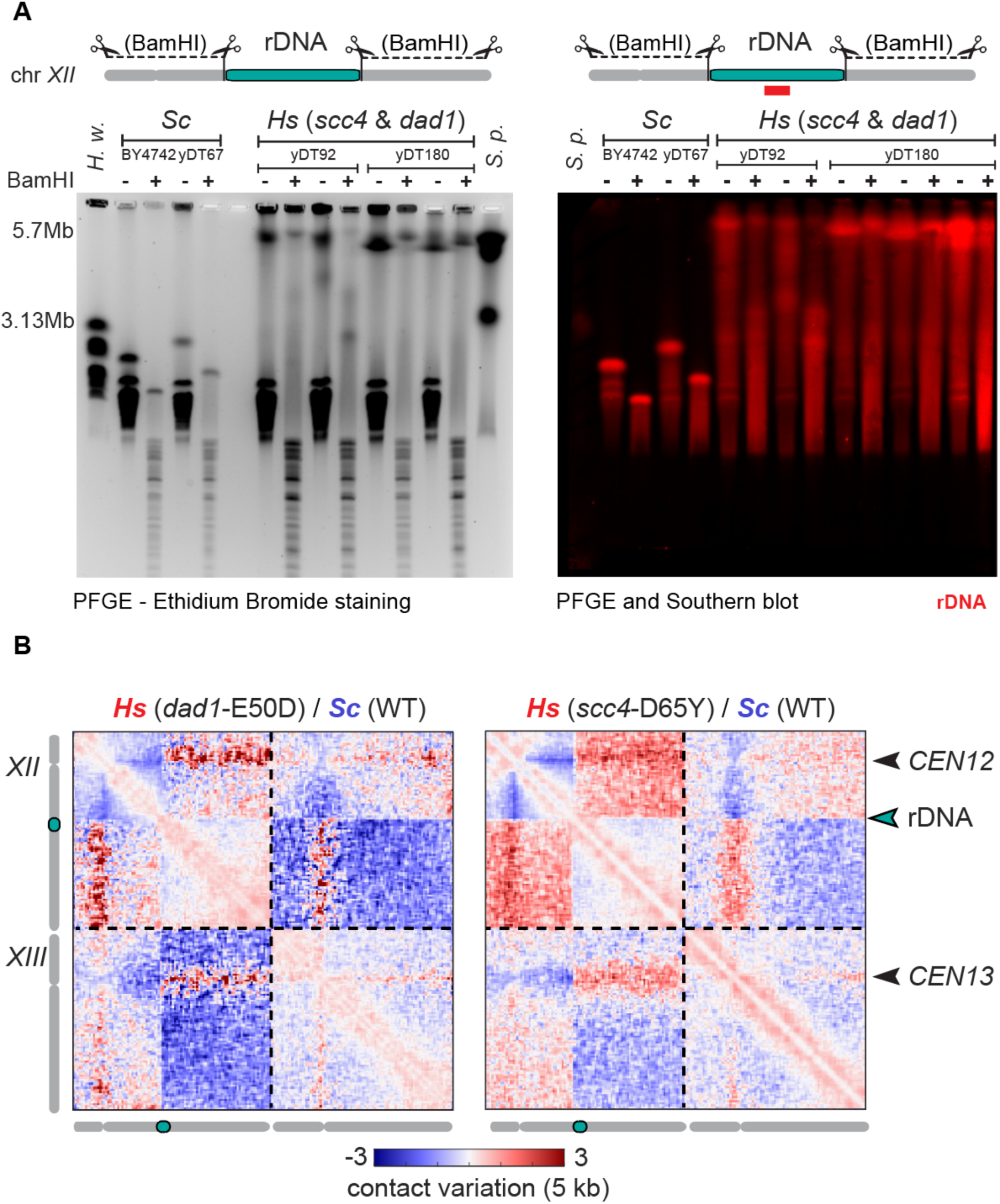
Histone humanization leads to the expansion of the rDNA array. (**A**) Estimated rDNA locus sizes (turquoise region on chromosome *XII*) in independent isolates of *Sc* (strains: BY4741, yDT67) and *Hs* (strains: yDT92, yDT180) yeasts. PFGE of chromosomes digested (+) or not (–) with BamHI (left panel) and the corresponding Southern blot (right panel) with an rDNA specific probe (red). PFGE ladders: *H. wingei* chromosomes (left) and *S. pombe* chromosomes (right). PFGE run specifications: *S. pombe* program for multi-megabase chromosome separation. (**B**) Contact map comparisons showing chromosomes *XII* and *XIII*. Log2-ratio maps of *Hs* vs. *Sc* strains: yDT180 *dad1*-E50D (left) and yDT92 *scc4*-D65Y (right). Arrowheads indicate the positions of the two centromeres and the rDNA locus. Color bar indicates contact variation between samples (log2 ratios 5 kb-binned).

**Figure S7, related to Main Figure 5.**
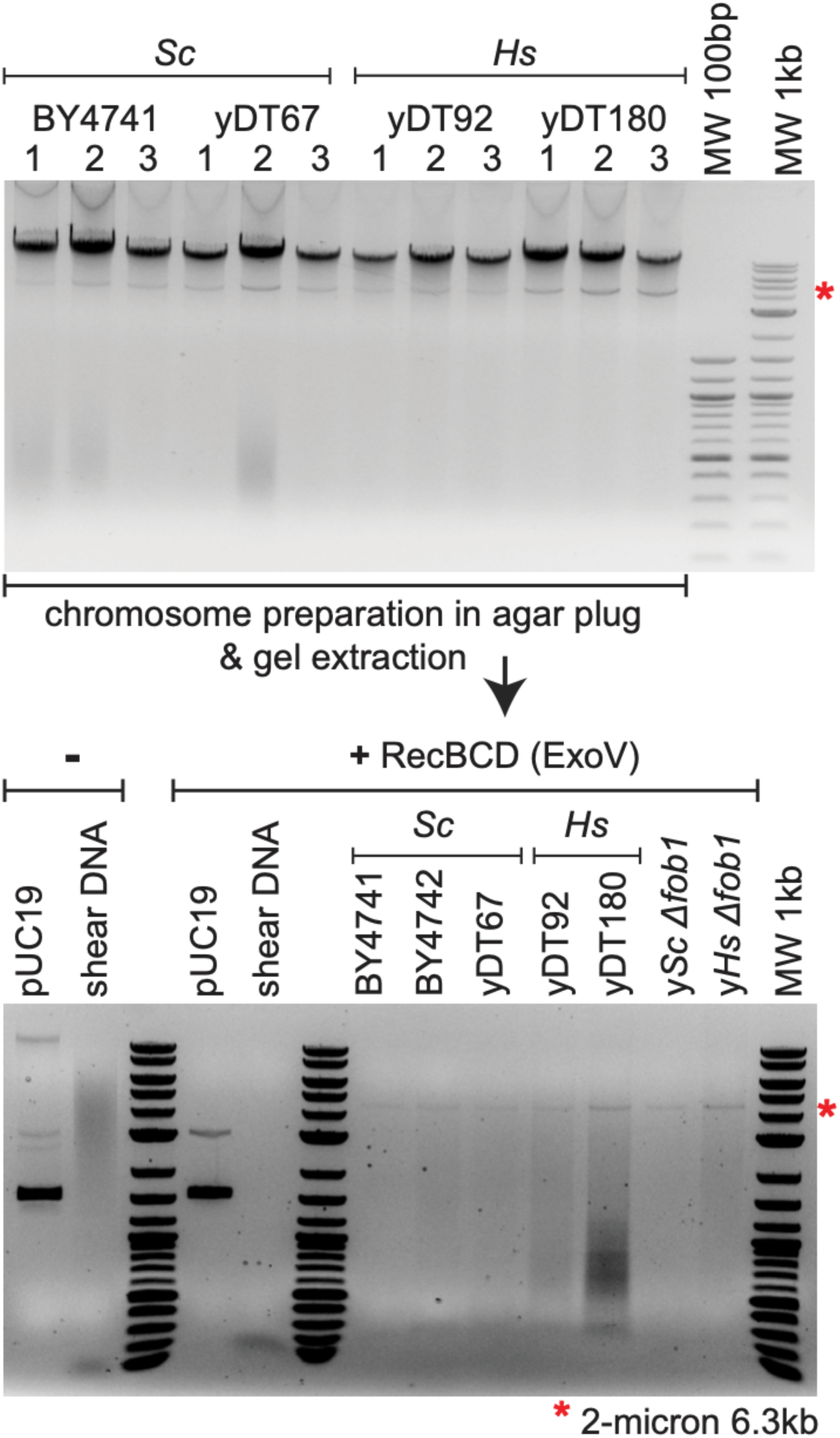
Histone humanization does not lead to extra-chromosomal rDNA circles. Agarose gels stained with ethidium bromide showing: (top panel) total genomic DNA extracted from *Sc* (strains: BY4741, yDT67) and *Hs* (strains: yDT92, yDT180) yeasts and (bottom panel) after RecBCD treatment. pUC19 circular plasmid and sheared DNA were used as controls. Note that strains with the *FOB1* gene deleted were also tested and represent negative controls for extra-chromosomal rDNA circles (ERCs) formation. Red (*) indicates 2-micron plasmid (∼40-60 copies/cell^127^).

**Figure S8, related to Main Figure 6.**
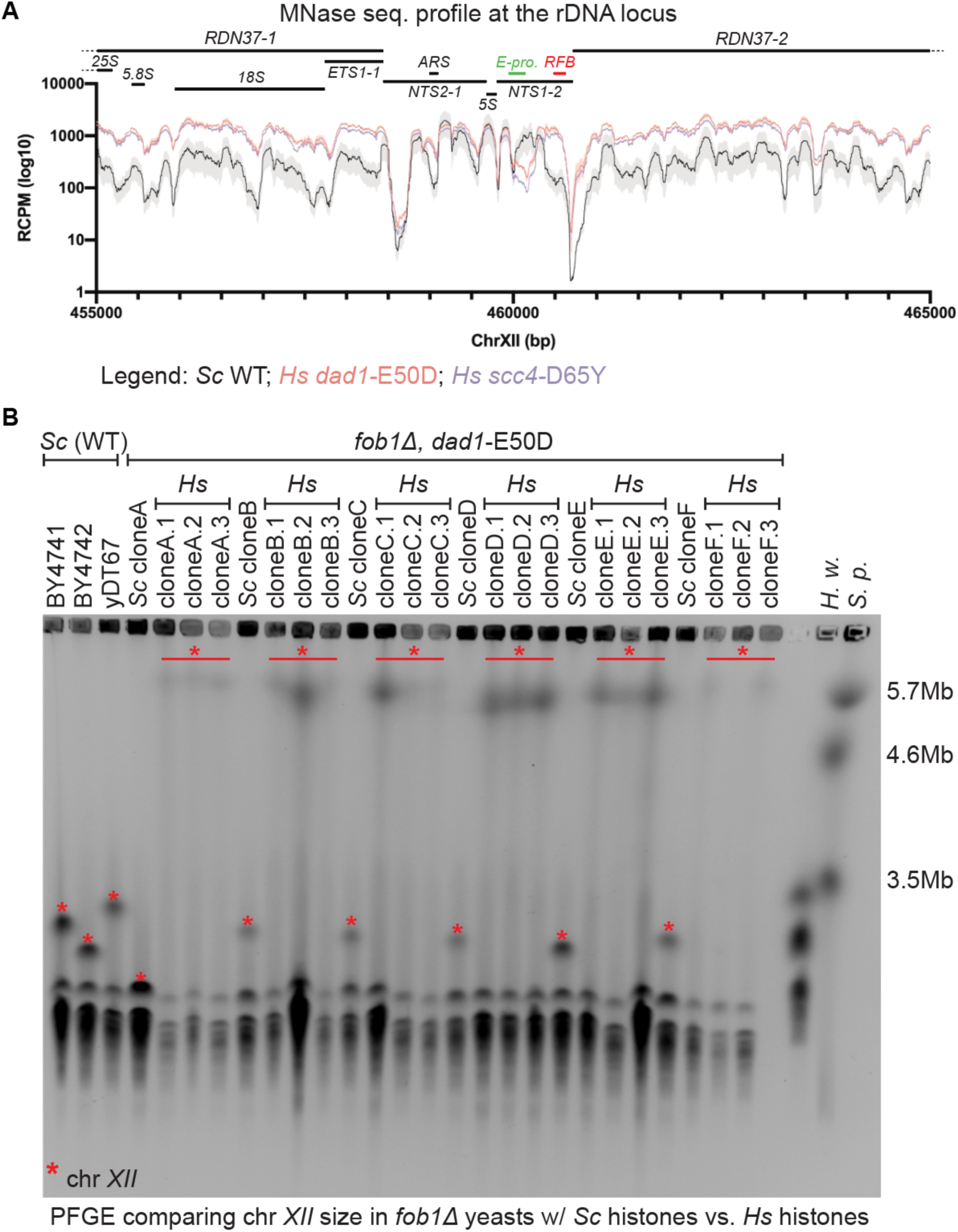
rDNA instability is independent of the replication fork block. (**A**) MNase-sequencing coverage profiles at the rDNA locus in *Sc* and *Hs* strains (re-analyzed data from Truong and Boeke^50^). (**B**) PFGE of yeast chromosomes in *fob1Δ* strains (Fob1, rDNA replication fork block-binding protein). *FOB1* was deleted in *Sc* (clones A to F; strains yMAH1242-12447) followed by histone humanization *Hs* (clones: A# to F#; yLS118-123). Each lane represents an independent isolated clone. PFGE ladders on the right: *H. wingei* and *S. pombe* chromosomes. (*) indicates chromosome *XII*. PFGE run specifications: *S. pombe* program for multi-megabase chromosome separation.

**Figure S9, related to Main Figure 6.**
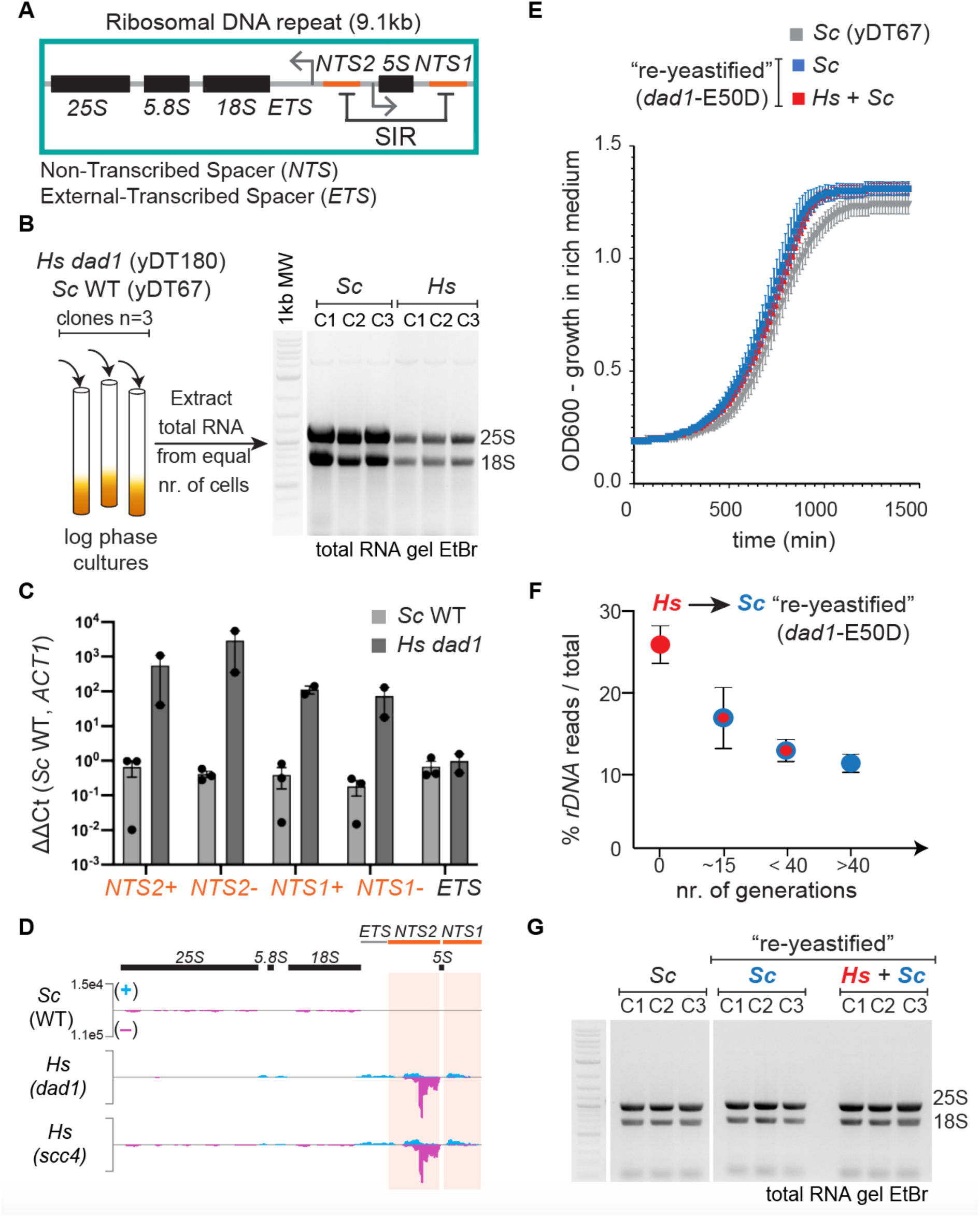
The epigenetic instability of the rDNA depends on human histones and is reversible. (**A**) Schematic showing the organization of a ribosomal DNA repeat unit with rRNA genes (*25S*, *18S*, *5.8S* and *5S*) and regulatory sequences (*NTS1* and *NTS2* silenced by SIR complex). (**B**) Diagram of RNA extractions from triplicates of *Sc* (yDT67) and *Hs* (yDT180 *dad1*-E50D) strains and agarose gel used for rRNA quantifications in Figure 6B. (**C**) RT-qPCR bar plot used to estimate changes in the transcription of the *NTS1/2* (“+” and “-“ DNA strands transcribed from the bidirectional E-promoter located in *NTS1*) and the rRNA precursor (*ETS*) relative to the control mRNA, *ACT1* (see Table S3). (**D**) Total RNA-sequencing coverage tracks at the rDNA unit in *Sc* and *Hs* strains (see Table S4). *y*-axis normalized to read counts per million. (**E**) Growth curves in rich media of the “re-yeastified” strains with *dad1-*E50D mutation (without *Hs* histones, *Sc*: yMAH753-755; *Hs* histones-maintained, *Hs* + *Sc*: yMAH756-758). (**F**) rDNA read count of the “re-yeastified” strains in Figure 5C. (**G**) RNA gel of the “re-yeastified” strains, as descried in panel E (*Sc*: yMAH753-755; *Hs* + *Sc*: yMAH756-758), relative to the wild-type *Sc* (yDT67) strain.

## Supplemental Excel Table Titles and Legends

**Table S1, related to Figure 1C. Mononucleosome surface area.** Summary of all measured mononucleosomes on yeast DNA with *S. cerevisiae* histones (*Sc* strain: BY4742) and human histones (*Hs* strains: yDT92 and yDT180).

**Table S2, related to Figure 5C. Estimating rDNA locus size.** (**A**) rDNA read counts in yeast strains with either wild-type histones (*Sc*) or human histones (*Hs*). (**B**) rDNA size after “re-yeastification” of the chromatin: swap *Hs* histones (pDT109) with the *Sc* histone plasmid (pDT105 or pDT139). (**C**) Expansion of the rDNA in independent histone-humanized yeast isolates carrying distinct humanization suppressor mutations.

**Table S3, related to Figure 6C and S9C. RT-qPCR measuring NTS transcription.** Raw Ct values of *NTS1*, *NTS2, ETS1* and *ACT1* transcripts in triplicates of yeast strains with *Sc* histones (strains: BY4741, yDT67) and *Hs* histones (yDT180).

**Table S4, related to Figure S9D. List of differentially expressed genes.** Combined RNA-sequencing data analysis from triplicates of yeast strains with *Sc* histones (yDT67) and *Hs* histones (yDT180). Re-analyzed data from Haase et al.^51^.

